# Kinase condensates enrich ATP and trigger autophosphorylation

**DOI:** 10.1101/2025.10.28.684909

**Authors:** Nicholas E. Lea, Lindsay B. Case

## Abstract

Kinase-mediated signal transduction regulates most cellular processes, and autophosphorylation is a common mechanism to activate kinases. Many kinases undergo phase separation to form condensates. Despite the central role of autophosphorylation in regulating kinase activity, how condensates impact kinase autophosphorylation has not been systematically studied. Using biochemical reconstitution and cellular studies, we find that phase separation can concentrate kinases to effectively trigger the *trans*-autophosphorylation of the tyrosine kinases FAK and Abl, as well as the serine/threonine kinase Mst2. Moreover, kinase condensates can create a chemical environment that enriches ATP, and positively charged intrinsically disordered regions are one feature that enrich ATP into condensates. Thus, kinase phase separation is a general mechanism to activate kinase signaling pathways by locally concentrating both kinases and ATP to trigger autophosphorylation.

## Introduction

Kinase activity plays a central role in signal transduction and regulates most cellular processes. High-fidelity signal transduction requires kinase activity to be tightly regulated, ensuring that substrate phosphorylation occurs at the correct time and place. Growing evidence suggests eukaryotic cells can regulate signal transduction by recruiting kinases to biomolecular condensates^1–3^. Condensates are membraneless cellular compartments that concentrate specific biomolecules and regulate diverse cellular processes^4–6^. Many condensates form through the phase separation of their constituent macromolecules, a thermodynamically spontaneous favorable process often driven by weak, multivalent protein interactions mediated by folded domains and intrinsically disordered regions (IDRs)^7^. Phase separation depends on molecular concentrations as well as solution conditions such as temperature, pH, salt, and molecular crowding. Phase separation results in two distinct phases, a “dense phase” highly concentrated in select molecules and a “dilute phase” that is much less concentrated^5^. In addition to concentrating and excluding specific macromolecules, condensates have emergent physical and chemical properties that can be important for their function^7–10^.

Many kinases directly undergo phase separation to drive condensate formation^6^. These kinase condensates activate signaling and regulate various cellular processes, including cell-volume homeostasis^11^, autophagy^12^, proliferation^13^, and survival^14^. In cancer, chromosomal rearrangements can result in oncogenic kinase fusion proteins that form cytoplasmic kinase condensates that hyperactivate signaling^14–17^. Kinase condensates can impact signaling through a variety of mechanisms. Concentrating substrates and kinases together within synthetic condensates is sufficient to increase substrate phosphorylation and broaden kinase specificity^20^. For Plk4 and PKA, the active conformation of the kinase is recruited to condensates^18,19^. In the Hippo signaling pathway, positive and negative regulators of the Hippo kinase cascade can both undergo phase separation. Positive regulators of Hippo signaling form condensates that locally concentrate kinases and activate their signaling, while negative regulators of Hippo signaling form condensates that concentrate kinases with phosphatases to inactivate signaling^13^.

The most common and conserved mechanism to activate eukaryotic kinases is autophosphorylation^21,22^. Many kinases perform this reaction *in trans*, a concentration-dependent process where one kinase molecule phosphorylates a second, identical kinase molecule. Despite the central role of autophosphorylation in regulating kinase activity, how condensates impact kinase autophosphorylation has not been systematically studied. Using biochemical reconstitution and cellular studies, we find that phase separation can concentrate kinases to effectively trigger the *trans*-autophosphorylation of cytoplasmic tyrosine and serine/threonine kinases. Moreover, positively charged IDRs of some kinases create a condensate chemical environment that concentrates adenosine triphosphate (ATP). Our data suggest that kinase phase separation is a general mechanism to activate kinase signaling pathways by locally concentrating both kinases and ATP, robustly triggering autophosphorylation.

## Results

### FAK condensates trigger autophosphorylation *in vitro*

Focal adhesion kinase (FAK) is a cytoplasmic tyrosine kinase that regulates integrin-dependent survival and migration^23^. FAK is typically autophosphorylated at focal adhesions, supramolecular assemblies that form at the plasma membrane when integrin receptors bind the extracellular matrix (ECM). FAK autophosphorylation creates a binding site for Src kinase, leading to phosphorylation of additional substrates^24^. We previously found that intermolecular interactions between FAK molecules enable the spontaneous phase separation of FAK to form micron-sized condensates at nanomolar concentrations^25^. Moreover, FAK phase separation contributes to initial focal adhesion formation in cells spreading on extracellular matrix^25^. FAK-dependent signaling requires FAK autophosphorylation on tyrosine 397 (Y397), which predominantly occurs *in trans*^26^. Thus, we used recombinant full-length FAK as a simple model system to study *trans*-autophosphorylation in the context of phase separation (**Figure S1A**). We found that both phosphorylated and dephosphorylated FAK undergo phase separation, although phosphorylated FAK has a slightly increased dense phase volume at similar concentrations (**Figure S1B-D**). Thus, tyrosine phosphorylation is not required for FAK phase separation and autophosphorylation does not dramatically change FAK phase separation propensity. To test if phase separation changes the rate of FAK autophosphorylation, we used perturbations that reduce or enhance FAK phase separation. As previously reported^25^, NaCl concentrations of 300 mM completely inhibit phase separation by preventing electrostatic interactions, a mutation of the FERM domain dimerization interface (W266A) decreases phase separation by reducing FAK dimerization, and the addition of 100 nM paxillin increases phase separation by introducing additional multivalent protein interactions (**Figure 1A-B**). Thus, if we maintain FAK at 1 µM concentration, the total condensate area significantly varies across these conditions (**Figure 1B**). We measured FAK autophosphorylation kinetics using a phosphospecific antibody for pY397 and found that conditions with increased condensate areas exhibit faster rates of autophosphorylation (**Figure 1C-D**). Next, we used a previously developed sedimentation assay^27^ to directly measure the dilute phase rate and indirectly assess the dense phase rate (**Figure 1E**). We observed that the initial rate of autophosphorylation is substantially slower in the dilute phase compared to the total solution and estimate that the dense phase accounts for >99% of the total solution activity (**Figure 1F-G**). Using quantitative microscopy, we found that the mEGFP-FAK concentration is 45 nM in the dilute phase and 106 µM in the dense phase (**Figure 1H-I**). Although FAK condensates account for most of the solution activity, they occupy <1% of the solution volume. For a given volume, we estimate that autophosphorylation rate is ∼90,000 times faster within condensates compared to the dilute phase (**Figure 1J**; see Supplementary Text for details).

**Figure 1.**
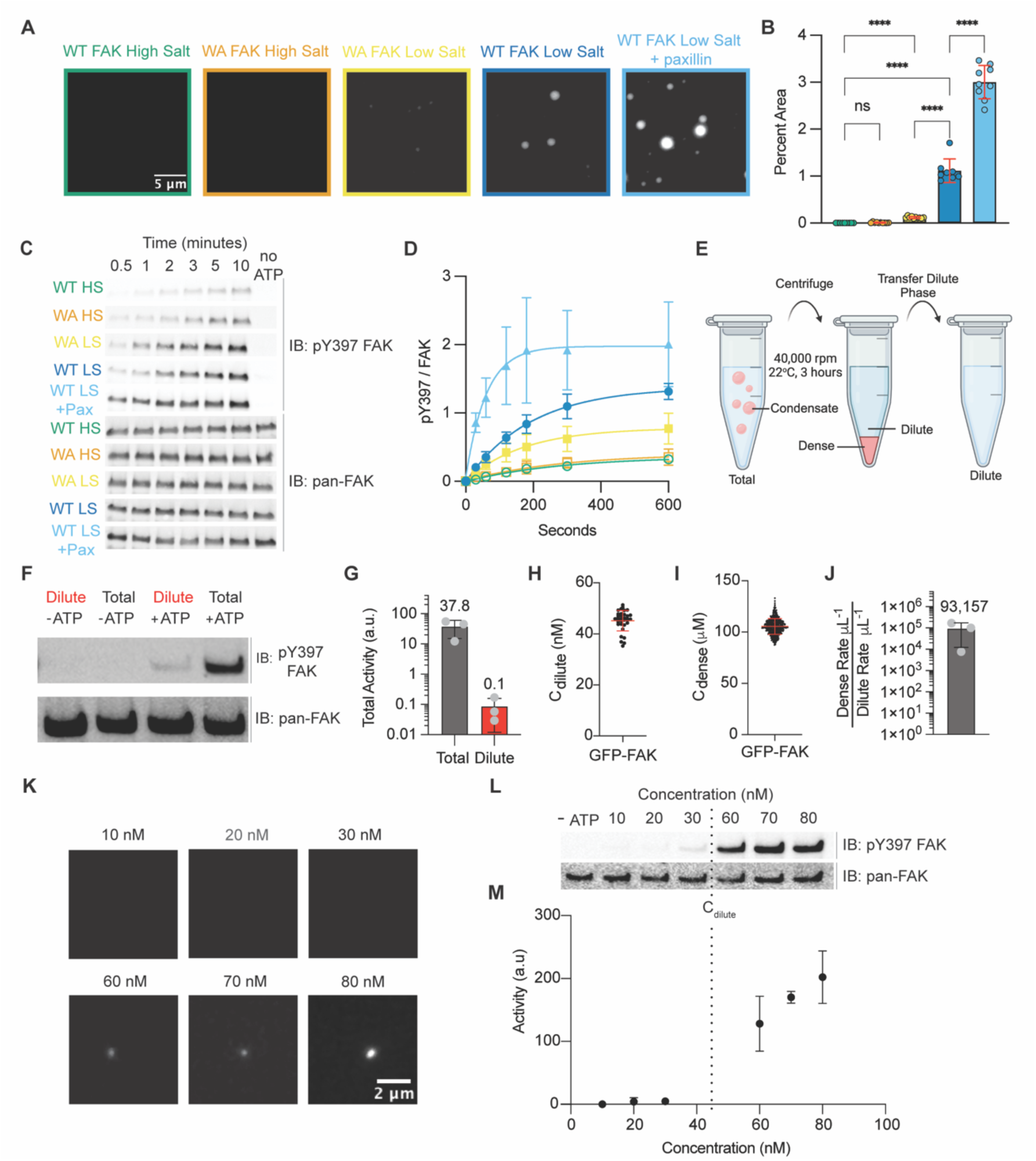
FAK condensates trigger autophosphorylation in vitro. (**A**) Images of 1 µM mEGFP-FAK with different buffers or mutations. Buffer: 25 mM HEPES pH 7.5, 50 (Low Salt) or 300 (High Salt) mM NaCl, 1% glycerol, 1 mM DTT. Paxillin is 100 nM. (**B**) Quantification of percent area occupied by condensates for conditions in (A). N > 7 images per condition. Significance was tested with Brown-Forsythe and Welch ANOVA with Dunnett’s T3 multiple comparison tests (* p<0.0332, ** p<0.0021, *** p<0.0002, **** p<0.0001). (**C**) Western blots of FAK autophosphorylation for each condition in (A). (**D**) Quantification of data in (C). N=3 replicates. (**E**) Diagram of separating condensates by centrifugation. (**F**) Western blot of total and dilute phase initial autophosphorylation rates at 30s. (**G**) Quantification of data in (F). N=3 replicates. (**H-I**) Measurements of mEGFP-FAK dilute phase concentration (H) and dense phase concentration (I) using fluorescence microscopy. N=34 images for C_dilute_ and N=502 droplets for C_dense_. (**J**) Volume-normalized fold rate enhancement of dense phase. (**K**) Images of mEGFP-FAK. Buffer matches Low Salt buffer in (A). (**L**) Western blot of FAK autophosphorylation. (**M**) Quantification of the data in (K) (N=3 replicates). Dotted line represents C_dilute_ value measured in (H). For all graphs, error bars are standard deviation and numbers above error bars are means.

Phase separation occurs at concentrations above the saturation concentration (C_sat_), which is also the concentration of protein in the dilute phase at equilibrium (C_dilute_)^28^. The mEGFP-FAK C_dilute_ is 45 nM, which we validated independently with quantitative microscopy (**Figure 1H**), quantitative western blots (**Figure S2A-C**), and titration (**Figure 1K**, **Figure S2D**). When we titrated mEGFP-FAK across its C_dilute_, we observed a 25-fold increase in autophosphorylation activity between concentrations just below and just above C_dilute_ (30 nM vs 60 nM) (**Figure 1K-M**). This increase in activity is more than expected from doubling the enzyme concentration, consistent with FAK condensates having drastically increased autophosphorylation activity. Additionally, we performed mass photometry on mEGFP-FAK and found there is no change in FAK oligomerization in the dilute phase across these concentrations (**Figure S2E**). We conclude that the rate of autophosphorylation significantly increases upon condensate formation due to increased activity in the dense phase. Together, these biochemical experiments demonstrate that phase separation is sufficient to trigger robust FAK *trans*-autophosphorylation on Y397 by locally increasing kinase concentration.

### FAK condensates trigger autophosphorylation in cells

We next sought to test if FAK phase separation is sufficient to trigger FAK autophosphorylation in cells. The cellular concentration of FAK is estimated to be 5-10 nM in fibroblasts^29^. Thus, endogenous FAK is maintained at concentrations below its C_sat_, and FAK phase separation is likely regulated by its local concentration at focal adhesions through protein and lipid interactions^25,30^. However, we hypothesized that overexpression of FAK to concentrations above its C_sat_ would promote spontaneous FAK phase separation in the cytoplasm. To test this, we transiently overexpressed mEGFP-FAK in mouse embryonic fibroblasts (MEFs). To prevent integrin-dependent FAK autophosphorylation, we plated cells on poly-D-Lysine coated plates for 30 minutes to allow cells to attach to the surface without integrin adhesion (**Figure 2A**). We found that cells expressing high levels of mEGFP-FAK contained cytoplasmic mEGFP-FAK puncta (**Figure 2B**). Although FAK can associate with and signal from endosomes^31^, these cytoplasmic FAK puncta did not co-localize with recycling (Rab11 positive) or early (EEA1 positive) endosomes (**Figure S3A-B**). Cytoplasmic mEGFP puncta exhibit moderate fluorescence recovery after photobleaching (FRAP) (t_1/2_ = 41.1 ± 14.5 seconds; 56 ± 13% recovery) (**Figure 2C**). Cytoplasmic puncta are slightly more dynamic than condensates *in vitro (*t_1/2_ = 64.5± 10.8 seconds; 35 ± 7% recovery) (**Figure S4A-B**) while less dynamic than focal adhesions at the plasma membrane (t_1/2_ = 9.8 ± 7.0 seconds; 57 ± 11% recovery) (**Figure S4C**). We conclude that FAK overexpression is sufficient to form dynamic cytoplasmic puncta distinct from endosomes.

**Figure 2.**
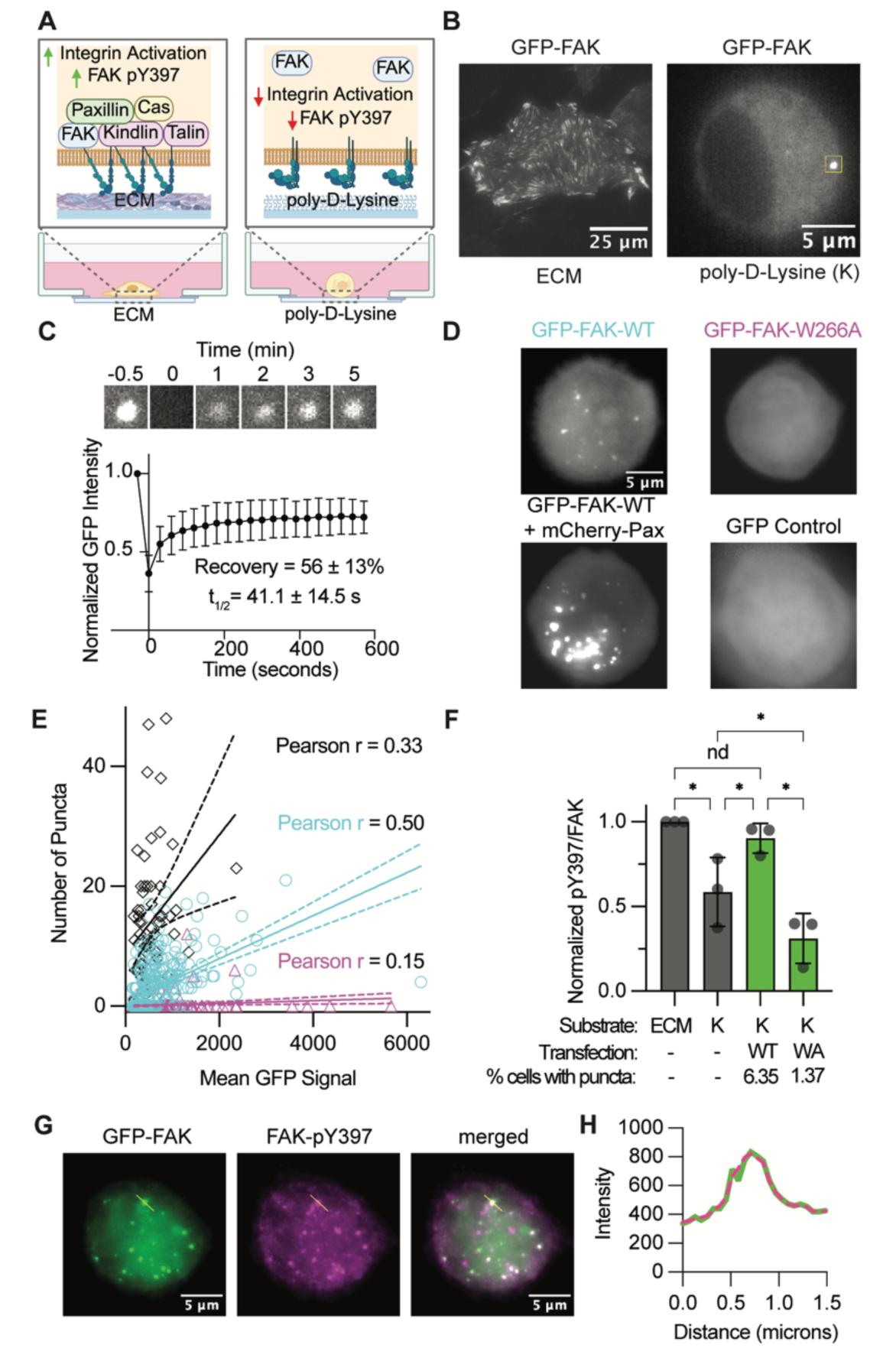
Cytoplasmic FAK condensates are sufficient to increase FAK autophosphorylation independent of integrin-based adhesion. (**A**) Diagram of MEFs plated on extracellular matrix (ECM) or poly-D-Lysine. (**B**) Representative images of mEGFP-FAK in MEFs. (**C**) Fluorescent Recovery After Photobleaching (FRAP) of the GFP-FAK puncta in (B) and FRAP analysis for N=5 replicates. (**D**) Representative maximum intensity projections of MEFs expressing GFP-tagged proteins. (**E**) Quantification of number of puncta in cells vs. total GFP signal (N> 60 cells per condition from two experimental replicates). Dotted lines represent 95% confidence intervals of linear regressions. (**F**) pY397/FAK ratios determined by western blot analysis of MEF lysates. K denotes cells plated on poly-D-Lysine for 30 minutes. ECM denotes cells grown on culture treated dishes for 24 hours. Green bars indicate analysis was performed on mEGFP-FAK and grey bars on endogenous FAK. Significance was tested with one-way ANOVA with correction for multiple comparisons using the two-stage linear step-up procedure of Benjamini, Krieger and Yekutieli (* p<0.05). (**G**) Immunostain images of MEFs expressing mEGFP-FAK and plated on poly-D-Lysine. (**H**) Linescan intensity of yellow line in (G). All data show mean ± standard deviation

To further investigate whether cytoplasmic FAK puncta are condensates formed through phase separation, we tested the same perturbations previously characterized *in vitro* (**Figure 1A**). Similar to *in vitro* experiments, mEGFP-FAK-W266A forms significantly fewer puncta than mEGFP-FAK-WT. Additionally, co-expressing mEGFP-FAK-WT or mEGFP-FAK-W266A with mCherry-Paxillin significantly increases the number of mEGFP puncta (**Figure 2D-E** and **Figure S5A-D**). The addition of mCherry-Paxillin also substantially increased the percent of transfected cells that contained puncta for both wild type and W266A FAK (**Figure S5E-F**). The number of puncta in a cell correlates positively with mEGFP-FAK expression levels, consistent with concentration-dependent phase separation (**Figure 2E**, **Figure S5D**). Occasionally, we observed large, non-spherical aggregates in cells with the highest mEGFP-FAK-WT expression levels (**Figure S5E**). We conclude that overexpressing FAK above the C_sat_ is sufficient to induce spontaneous condensate formation in the cytoplasm through the same interactions that drive FAK phase separation *in vitro*.

We next tested if cytoplasmic FAK condensates are sufficient to promote autophosphorylation in cells. As previously reported, cells plated on poly-D-Lysine for 30 minutes have reduced pY397/total FAK ratios compared to cells plated on tissue culture plates for 24 hours, enough time to secrete ECM and form integrin-based adhesions (**Figure 2F**)^32^. However, overexpressing mEGFP-FAK-WT in cells at levels that induce cytoplasmic condensates in 6.4% of cells is sufficient to rescue autophosphorylation on poly-Lysine (**Figure 2F**), consistent with cytoplasmic FAK condensates increasing autophosphorylation in cells. In contrast, overexpressing mEGFP-FAK-W266A, which rarely forms condensates, did not rescue autophosphorylation (**Figure 2F**). Moreover, expressing mEGFP-FAK-WT at lower levels (∼1.5 fold above endogenous) that do not induce cytoplasmic condensates does not rescue autophosphorylation, indicating that FAK overexpression does not increase cellular pY397/FAK ratios in the absence of condensates. (**Figure S6**). Although cytoplasmic FAK condensates promote FAK autophosphorylation, we did not observe significant increases in downstream paxillin or p130Cas phosphorylation 30 minutes after plating on Poly-Lysine (**Figure S7**). Since Src is required for downstream FAK signaling^24^, we speculate that Src-dependent phosphorylation is not rescued in condensates in this time frame. Finally, we performed immunostaining and found that cytoplasmic FAK condensates contain pY397 FAK (**Figure 2G-H**). We conclude that cytoplasmic FAK condensates are sufficient to trigger FAK autophosphorylation even in the absence of upstream integrin activation.

### Abl and Mst2:Sav1 condensates trigger autophosphorylation in vitro

Based on our observations of FAK condensates, we hypothesized that phase separation would be sufficient to trigger the *trans*-autophosphorylation of any kinase. Using phase separation predictors developed with machine learning ^33–35^, we estimate that 13-44% of human kinases have a strong likelihood of undergoing phase separation (**Figure 3A**; **Data S1**). To test if phase separation is a general mechanism to trigger autophosphorylation, we characterized the phase separation and autophosphorylation of two additional kinases predicted to phase separate, Abl and Mst2, using biochemical approaches. Abl is a tyrosine kinase and proto-oncogene with a 460 amino acid IDR^36^. Mst2 is a serine/threonine kinase in the Hippo signaling pathway that phase separates when combined with the intrinsically disordered adaptor protein Sav1^13,37^. Both kinases are predicted to phase separate and are activated by autophosphorylation *in trans*^36,38^. Consistent with the prediction, we found that Abl undergoes concentration dependent phase separation on its own. Sav1 also undergoes concentration dependent phase separation alone and enriches Mst2, as previously reported (**Figure 3B-C**, **Figure S8**)^37^. The exchange of protein between condensates and the dilute phase was much slower for Abl and Mst2-Sav1 condensates than for FAK condensates (**Figure 3D**). However, Hippo pathway activation has been shown to require solid-like condensates, suggesting less dynamic kinase condensates can activate signaling pathways in cells^8^. We next used in vitro kinase assays to determine the phosphorylation activity in solutions containing condensates. We first confirmed the dephosphorylation and activity of purified Abl and Mst2 with phospho-specific antibodies to Y226 (Abl) and T180 (Mst2) (**Figure S9** A-B). We then used an ADPGlo luciferase assay for high-sensitivity and robust quantitation of phosphorylation rates in the total solution and dilute phase. Similar to FAK, the total solution of Abl and Mst2 showed fast initial rates of phosphorylation, while almost no phosphorylation occurred after removing condensates from the dilute phase (**Figure 3E**, **Figure S9C**). We used phospho-proteomics analysis to determine the specific residues that were modified in these experiments (**Figure** S9D-G). While the only phospho-tyrosine detected on FAK was the canonical Y397 site^32^, Abl and Mst2 phosphorylated additional residues beyond their canonical autophosphorylation sites^36,38^. This observation suggests that condensates expand the specificity of autophosphorylation for some kinases, consistent with previous studies demonstrating that synthetic kinase condensates can phosphorylate unexpected peptides^20^. We conclude that phase separation is a general mechanism that can trigger *trans*-autophosphorylation of both tyrosine and serine/threonine kinases.

**Figure 3.**
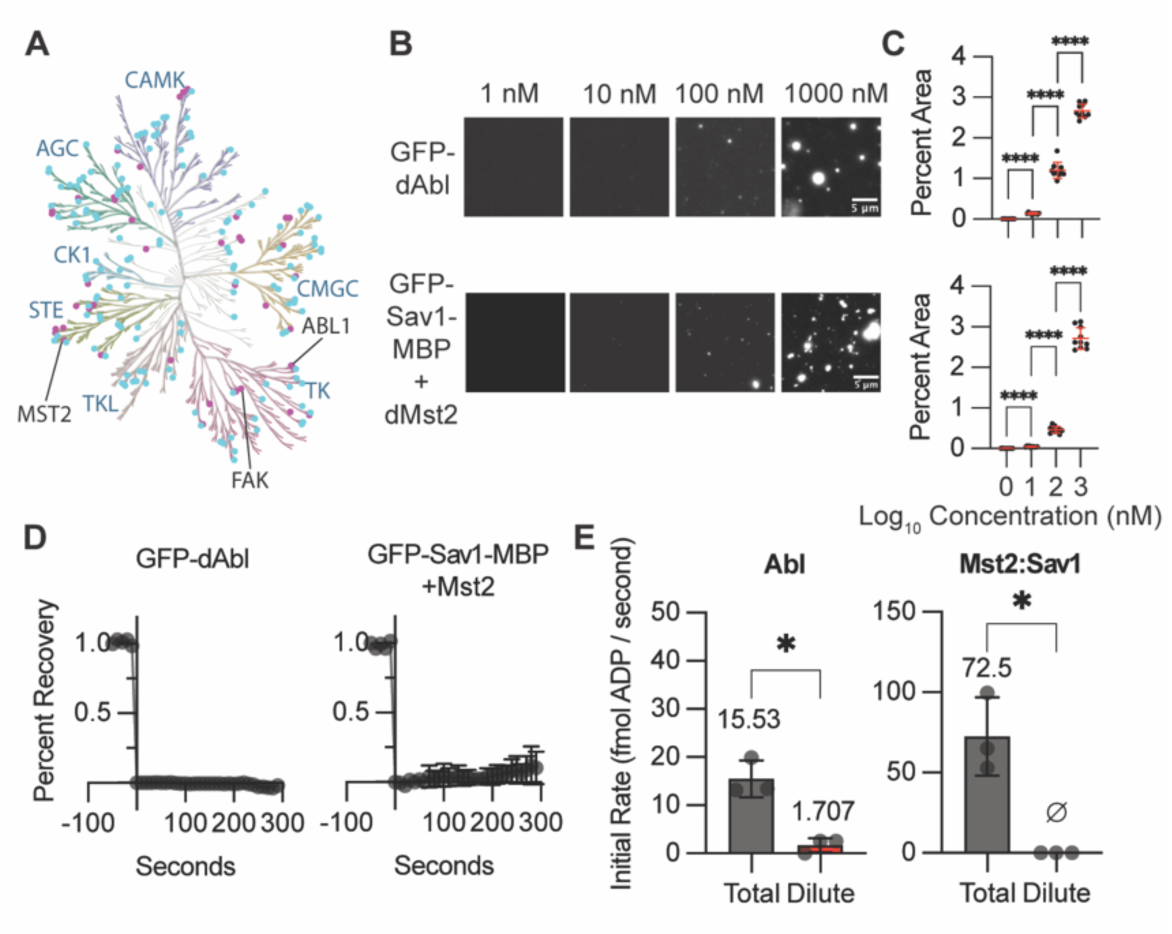
Abl and Mst2:Sav1 condensates trigger autophosphorylation in vitro. (**A**) Homology dendrogram of the human kinome. Kinases predicted to phase separate are denoted by colored dots (at least one predictor = cyan; all three predictors = magenta). (**B**) Images of mEGFP-dAbl and mEGFP-Sav1-MBP + dMst2. mEGFP-Sav1-MBP and dMst2 are kept at equimolar concentrations. Abl buffer is 25 mM HEPES pH 7.5, 100 mM NaCl, 5% PEG8000, 1% glycerol (v/v), 1 mM DTT. Sav1:Mst2 buffer is 25 mM HEPES pH 7.5, 100 mM NaCl, 10% PEG8000, 1 mM DTT. (**C**) Quantification of percent area occupied by condensates for conditions in (A). N = 10 images per condition. Significance was tested with Brown-Forsythe and Welch ANOVA with Dunnett’s T3 multiple comparison tests. (**D**) Fluorescence recovery after photobleaching (FRAP) analysis of mEGFP-dAbl and mEGFP-Sav1-MBP + dMst2 condensates. mEGFP-dAbl is 1,000 nM. mEGFP-Sav1-MBP and dMst2 are 40 nM. Buffers same as (B). N=5 replicates. (**E**) Initial rates of kinase phosphorylation assays. Buffers same as (B), except supplemented with 0.5 mM MgCl_2_ and initiated by addition of 1 μM ATP. N=3 replicates. Number above error bars is the mean. ø symbol denotes activity was undetectable. Significance was tested with unpaired t-test with Welch’s correction. For all graphs, error bars represent standard deviation. * p<0.0332, ** p<0.0021, *** p<0.0002, **** p<0.0001.

### Kinase condensates enrich ATP independent of active site binding

In addition to concentrating proteins, condensates create unique environments with emergent chemical and physical properties^7^. For example, condensates create distinct chemical environments that can enrich or exclude specific small molecules and metabolites^39,40^. Since ATP is a critical substrate for kinases, we sought to determine the extent to which ATP is enriched or excluded from reconstituted kinase condensates. At micromolar concentrations, a fluorescent analog of ATP (Alexa647-ATP) is enriched within FAK and Abl condensates, but neither enriched nor excluded from Mst2 condensates (**Figure 4A**). The free Alexa647 dye does not show enrichment or exclusion from these condensates and BODIPY-ATP-y-S is similarly enriched in FAK condensates, indicating that ATP is likely driving this enrichment (**Figure S10A-B**). Since fluorescent dyes can alter the chemical properties of nucleotides and often exhibit different fluorescent properties depending on chemical environment, we developed a luciferase-based assay to directly quantify unmodified nucleotide concentrations in the dense and dilute phases (**Figure 4B**). We validated this assay by comparing to spectrophotometric measurements of nucleotide concentrations in poly-lysine condensates (**Figure S11**)^41^. When we added 50 µM ATP to experiments with kinase condensates, the dense phase ATP concentrations were significantly higher than the dilute phase for FAK and Abl condensates (194 µM and 280 µM, respectively) while not significantly different for Mst2:Sav1 condensates (72 µM; **Figure 4C**). Magnesium was omitted from these experiments to prevent binding of ATP to the kinase active site and to prevent autophosphorylation activity in the samples^42^. We also found that ADP and GTP were significantly enriched in the dense phase of FAK condensates (**Figure 4 D-E**). Since GTP has undetectable binding to the FAK active site^43^, we conclude that kinase condensates can enrich nucleotides independent of high affinity stereospecific binding to the active site.

**Figure 4.**
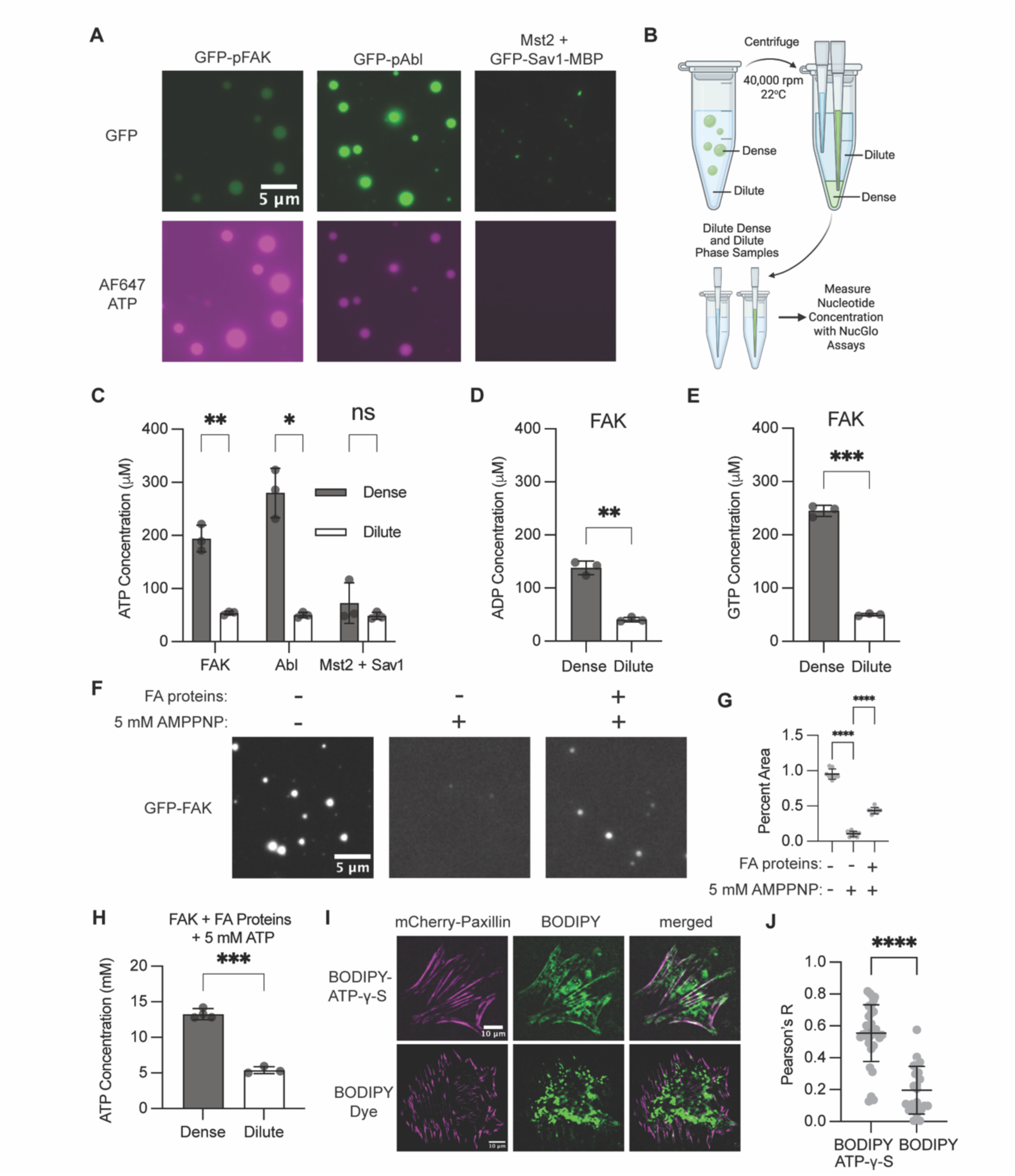
FAK and Abl condensates enrich ATP independent of active site binding. (**A**) Images of 1 μM mEGFP-FAK, 1 μM mEGFP-Abl, or 100 nM mEGFP-Sav1-MBP + 100 nM Mst2 with 2.5 μM AF647-ATP. (**B**) Diagram of sedimentation-luciferase assays for nucleotide measurements. **(C-E)** Measurements of ATP (C), ADP (D), and GTP (E) concentrations in the dense and dilute phase with 50 μM total nucleotide. (**F**) Images of 1 μM mEGFP-FAK. “FA proteins” denotes the addition of the purified focal adhesion proteins paxillin, p130Cas, Nck and N-WASP at 1 μM. (**G**) Quantification of percent area occupied by condensates in (F). N=8 images. Significance tested with Brown-Forsythe and Welch ANOVA with Dunnett’s T3 multiple comparison tests. (**H**) Measurements of ATP concentration in the dense and dilute phase with 5 mM total ATP. (**I**) Representative TIRF microscopy images of MEFs. (**J**) Pearson’s correlation coefficient between focal adhesions (mCherry-paxillin marker) and BODIPY-ATP-γ-S or BODIPY dye alone (N=35 and 23 cells respectively). For (C), (D), (E), (H) and (J) significance was tested with unpaired t-tests with Welch’s correction. For all sedimentation-luciferase assays, N=3 replicates. For all in vitro experiments, buffer conditions match those in Figure 1A (FAK) and Figure 3B (Abl and Sav1+Mst2). For all graphs error bars represent standard deviation and * p<0.0332, ** p<0.0021, *** p<0.0002, **** p<0.0001

Next, we wanted to determine if kinase condensates could enrich ATP at physiological nucleotide concentrations. In cells, ATP is complexed with magnesium and is maintained at millimolar concentrations in 20-1,000 fold excess of ADP^44^. However, millimolar concentrations of ATP can also reduce or enhance phase separation depending on the context^41,45,46^. To determine if kinase phase separation was sensitive to ATP independent of kinase ATPase activity, we added 5 mM of AMPPNP, a non-hydrolyzable ATP analog, to *in vitro* condensate assays (**Figure S12**). We found that Mst2 was unaffected by AMPPNP, Abl formed more condensates, and FAK formed very few condensates (**Figure S12A-B**). Combining FAK with additional adaptor proteins to form more physiological focal adhesion condensates^25^ partially rescued condensate formation in millimolar AMPPNP (**Figure 4F-G**). Next, we used the luciferase assay to measure ATP enrichment in focal adhesion condensates. When we added 5 mM ATP to focal adhesion condensates, the dense phase contained 13.3 mM ATP (**Figure 4H**). The ATP concentration in the dense phase was not altered by including 5 µM ADP in the experiment (**Figure S13**). We also used microscopy to assess ATP enrichment in kinase condensates at millimolar concentrations (**Figure S14**). When we used buffer containing 1 mM AMPPNP and 10 μM Alexa647-ATP, we found that focal adhesion condensates and Abl condensates enriched ATP even with 1mM MgCl_2_ and 50 μM ADP (**Figure S14A**). Mst2 condensates did not enrich ATP in any conditions. When we used 1 mM ATP instead of AMPPNP we saw similar Alexa647-ATP enrichment in focal adhesion condensates across all conditions, but less enrichment in Abl condensates with 1mM MgCl_2._ Including 1 mM ATP-Mg supports rapid autophosphorylation of both Abl and Mst2 (**Figure S9 D-G**), and this data suggests that the negative charge added by autophosphorylation reduces ATP enrichment in Abl condensates. Together, these data demonstrate that focal adhesion condensates and Abl condensates enrich ATP at physiological nucleotide concentrations.

To further explore whether ATP enrichment into kinase condensates could be physiologically relevant, we tested the ability of cellular focal adhesions to concentrate BODIPY-ATP-y-S. Using a previously developed assay to introduce molecules into cells^47^, we gently permeabilized cells and incubated with BODIPY-ATP-y-S or BODIPY. We found that BODIPY-ATP-y-S was non-uniform in its localization in the cytoplasm and nucleus and appeared to enrich in focal adhesions (**Figure 4I**). Compared to BODIPY, BODIPY-ATP-y-S was significantly more colocalized with the focal adhesion marker paxillin (**Figure 4I-J**). We conclude that ATP can enrich within both reconstituted focal adhesion condensates and cellular focal adhesions.

### Positive charge is sufficient to drive ATP enrichment into IDR condensates

FAK and Abl condensates both enrich ATP, and both kinases contain IDRs (**Figure 5A**). Previous studies have found that ATP is highly enriched in condensates formed from positively charged IDRs ^41,46,48^. Thus, we hypothesized that positive charge within IDRs could be important for ATP partitioning into kinase condensates. FAK has a negatively charged ∼200 amino acid IDR (net charge per residue = - 0.02), while Abl has a positively charged ∼460 amino acid IDR (net charge per residue = + 0.05) (**Figure 5A**). Both IDRs form condensates in the presence of crowding agent (20% PEG), and but only Abl IDR condensates enrich Alexa647-ATP (**Figure 5B**).When we added 50 µM ATP, the Abl IDR dense phase enriches ATP 20-fold (773 µM in the dense phase; 32 µM in the dilute phase), but the FAK IDR dense phase does not significantly enrich ATP (59 µM in the dense phase; 45 µM in the dilute phase) (**Figure 5C**). To further test if positive charge is sufficient to drive ATP enrichment, we investigated the C-terminal IDR of DDX21. DDX21 is a DEAD-box ATPase with a well-characterized positively charged C-terminal IDR (net charge per residue = + 0.22) that phase separates ^49^. The DDX21 IDR dense phase enriches ATP 20-fold (931 µM in the dense phase; 37 µM in the dilute phase), and neutralizing the IDR charge is sufficient to abolish ATP enrichment (54 µM in the dense phase; 58 µM in the dilute phase; **Figure 5B-C**). Additionally, Abl and DDX21 IDR condensates enrich ATP when we added 3 mM ATP (5.8-fold ATP enrichment for Abl IDR; 15-fold ATP enrichment for DDX21 IDR; 2.6-fold ATP enrichment for DDX21 R/S IDR; **Figure S15A-B**). We conclude that positive charge is sufficient to enrich ATP within IDR condensates and that charge is an emergent condensate property that influences nucleotide partitioning.

**Figure 5.**
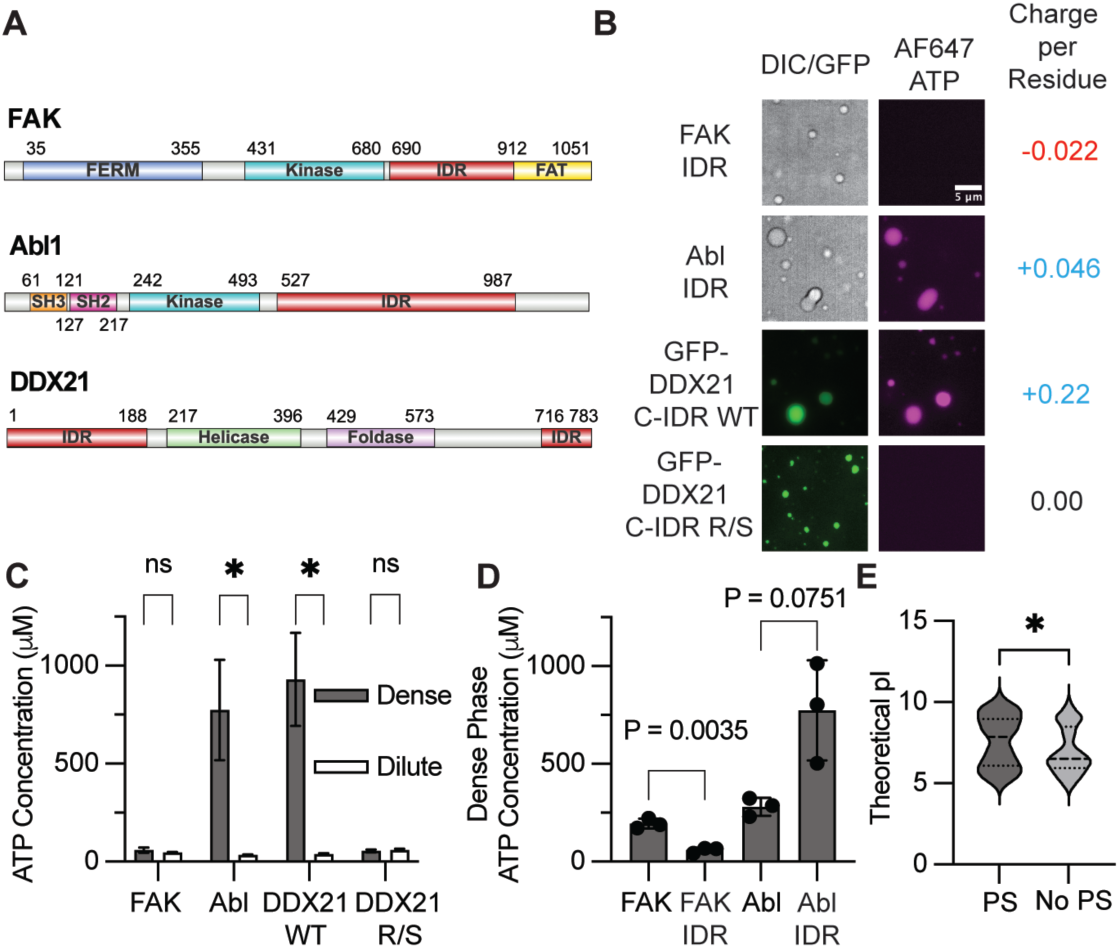
Positive charge is sufficient to enrich ATP in IDR condensates. (**A**) Diagram of FAK, Abl1 and DDX21 proteins indicating intrinsically disordered regions (IDR). (**B**) Images of IDR condensates with AF647-ATP. Buffer conditions are 20 μM protein, 25 mM HEPES pH 7.5, 100 mM NaCl, 20% PEG8000, 1 mM DTT and 2.5 μM AF647-ATP. (**C**) Measurements of ATP concentrations in dense and dilute phase from sedimentation-luciferase assays. Buffer conditions are identical to those in (B) except with 50 μM ATP instead of AF647 ATP (N=3 replicates). (**D**) Comparison of dense phase ATP concentrations between full length protein condensates and IDR condensates. Data is replotted from (C) and Figure 4C. For (C-D), error bars denote standard deviation and statistical tests are unpaired T-tests with Welch correction. (**E**) Distribution of theoretical isoelectric points (pI) of human protein kinases binned by those predicted (PS, N=68) and those not predicted (No PS, N=94) to phase separate. Statistical test is a Mann-Whitney test. Dotted lines represent quartiles. * p<0.0332, ** p<0.0021, *** p<0.0002, **** p<0.0001.

### Folded Domains Contribute to ATP Enrichment in Kinase Condensates

Over 80% of human kinases contain both folded domains and IDRs^50^. Although FAK has a negatively charged IDR that is not sufficient to enrich ATP, the full-length FAK protein forms condensates that enrich ATP (**Figure 5D**). Additionally, full-length Abl forms condensates that enrich less ATP than the Abl IDR condensates (**Figure 5D**). Thus, folded domains also contribute to ATP enrichment into condensates. Electrostatic interactions between ATP and folded domains and physicochemical properties beyond charge could contribute to ATP enrichment in condensates ^39,51^. Since kinase condensates enrich ATP at least in part due to positively charged residues, we computed the theoretical isoelectric point of human kinases. We found that kinases predicted to phase separate have significantly higher isoelectric points compared with kinases predicted to not phase separate (**Figure 5E**), consistent with a functional role for positive charge within kinase condensates. Together, our experimental data demonstrate that substrates can be enriched through the emergent chemical properties of enzyme condensates.

## Discussion

Hundreds of human kinases are likely activated through *trans-*autophosphorylation^22^, and our data demonstrate that phase separation is a robust mechanism to activate these kinases. Using in vitro kinase assays, we found that condensates elicit switch-like, rapid, and robust autophosphorylation of cytosolic tyrosine (FAK and Abl) and serine/threonine (Mst2) kinases. In cells, cytoplasmic FAK condensates are sufficient to trigger FAK autophosphorylation decoupled from upstream integrin receptor activation. FAK and Abl condensates also enrich ATP independent of stereospecific active site binding. While positively charged IDRs are sufficient to enrich ATP into condensates, folded domains also contribute to ATP enrichment in kinase condensates. Together, our data demonstrate that kinase phase separation is a general mechanism to activate kinase signaling pathways by locally concentrating both kinases and ATP to trigger *trans*-autophosphorylation.

Kinase phase separation is dependent on both kinase concentration and solution conditions such as temperature, pH, salt, and molecular crowding^28^. Our findings suggest that kinase phase separation could directly activate numerous kinase signaling pathways in response to changes in cellular conditions that promote phase separation^6^. Small changes in cellular conditions could promote kinase phase separation leading directly to autophosphorylation and activation of signaling. For example, WNK is a kinase that phase separates upon molecular crowding during hypertonic stress, and WNK condensate formation correlates with signaling activation^11^. Our data also reveal how kinase condensates can potentially dysregulate signaling pathways. We found that kinase overexpression is sufficient to form cytoplasmic condensates that decouple autophosphorylation from upstream receptor activation. FAK and Abl are both overexpressed in certain cancers, and kinase overexpression could trigger aberrant condensate formation to constitutively activate kinases^52,53^. Thus, gene fusions that create chimeric fusion kinases^14–17^ and overexpression of endogenous kinases may both result in kinase condensates that dysregulate signaling and decouple kinase activation from normal stimuli.

Condensates do not simply concentrate proteins but also create local chemical environments distinct from the dilute phase. We found that the emergent physicochemical properties of kinase condensates can drive nucleotide enrichment independent of stereospecific active site binding. This enrichment is driven, in part, by positively charged IDRs, although folded domains also contribute. In addition to influencing ATP enrichment, positively charged IDRs can also promote phase separation through interactions with ATP^41,46^. Kinases predicted to phase separate tend to have higher isoelectric points, consistent with a functional role for positive charge within kinase condensates. Moreover, extensive phosphorylation can dramatically reduce positive charge and would thus be expected to reduce ATP partitioning. This suggests that the dynamic modulation of protein charge through phosphorylation can negatively feedback to tune ATP enrichment and kinase activity inside condensates.

When we add millimolar concentrations of ATP to in vitro assays, condensates can enrich ATP 2-15-fold higher. Since most kinases have micromolar K_M_ for ATP, would this ATP enrichment impact autophosphorylation? The thermodynamic favorability of phosphorylation depends not on the ATP concentration but on the ratio of ATP to ADP. One important function of traditional membrane-bound organelles is to regulate ATP:ADP ratios to control biochemistry. ADP/ATP transport through the inner mitochondrial membrane maintains low ATP:ADP ratios inside mitochondria to thermodynamically drive ATP synthesis from ADP, while simultaneously maintaining high ATP:ADP ratios in the cytoplasm to favor ATP consuming processes. Our data demonstrate that ADP is less enriched than ATP in kinase condensates, likely because ATP is more negatively charged. This suggests that the ATP:ADP ratios in kinase condensates would be higher than in the surrounding dilute phase or cytoplasm. An active kinase will rapidly deplete ATP and generate ADP, which thermodynamically opposes further phosphorylation. The ability of kinase condensates to maintain high ATP:ADP ratios may counteract the consumption of ATP to sustain high rates of phosphorylation inside active kinase condensates. Maintaining a high ATP:ADP ratio in condensates could help drive phosphorylation in normal cell processes and potentially sustain kinase activity in cellular conditions where cytoplasmic ATP:ADP ratios are lower than normal, such as in cancer cells^55^.

## Acknowledgments

We thank Eliezer Calo (MIT) for providing DDX21 IDR DNA and the Super PiggyBac Transposase plasmid, Seychelle Vos (MIT) for providing insect cells and guidance with Baculovirus expression systems, Barbara Imperiali (MIT) for use of a BioTek hybrid reader, Michael Yaffe (MIT) and Rick Young (MIT) for helpful discussions about experimental controls, the Koch Institute Biopolymers & Proteomics Facility for phospho-mass spectrometry analysis, the Biophysical Instrumentation Facility at MIT for mass photometry assistance, the W.M. Keck Microscopy Facility at the Whitehead Institute for Spinning Disk Microscopy assistance, the Flow Cytometry Core Facility at the Whitehead Institute for cell sorting, and the Preclinical Modeling, Imaging & Testing Core at the Koch Institute for mycoplasma testing.

## Author contributions

Conceptualization: LBC, NEL

Methodology: LBC, NEL

Investigation: NEL

Visualization: NEL

Funding acquisition: LBC

Project administration: LBC

Supervision: LBC

Writing – original draft: NEL, LBC

Writing – review & editing: NEL, LBC

## Data and materials availability

All analyzed data are available in the main text or the supplementary materials. Raw microscopy images will be deposited to an online repository prior to publication. Unpublished plasmids will be deposited to Addgene prior to publication.

## References

1. Zhang, H., Cao, X., Tang, M., Zhong, G., Si, Y., Li, H., Zhu, F., Liao, Q., Li, L., Zhao, J., et al. (2021). A subcellular map of the human kinome. eLife 10, e64943. 10.7554/eLife.64943.

2. Li, P., Chen, P., Qi, F., Shi, J., Zhu, W., Li, J., Zhang, P., Xie, H., Li, L., Lei, M., et al. (2024). High-throughput and proteome-wide discovery of endogenous biomolecular condensates. Nat. Chem. 16, 1101–1112. 10.1038/s41557-024-01485-1.

3. López-Palacios, T.P., and Andersen, J.L. (2023). Kinase regulation by liquid–liquid phase separation. Trends in Cell Biology 33, 649–666. 10.1016/j.tcb.2022.11.009.

4. Banani, S.F., Lee, H.O., Hyman, A.A., and Rosen, M.K. (2017). Biomolecular condensates: organizers of cellular biochemistry. Nat Rev Mol Cell Biol 18, 285–298. 10.1038/nrm.2017.7.

5. Shin, Y., and Brangwynne, C.P. (2017). Liquid phase condensation in cell physiology and disease. Science 357, eaaf4382. 10.1126/science.aaf4382.

6. Cheng, X., and Case, L.B. (2023). Phase separation in chemical and mechanical signal transduction. Curr Opin Cell Biol 85, 102243. 10.1016/j.ceb.2023.102243.

7. Holehouse, A.S., and Alberti, S. (2025). Molecular determinants of condensate composition. Molecular Cell 85, 290–308. 10.1016/j.molcel.2024.12.021.

8. Guo, P., Li, B., Dong, W., Zhou, H., Wang, L., Su, T., Carl, C., Zheng, Y., Hong, Y., Deng, H., et al. (2024). PI4P-mediated solid-like Merlin condensates orchestrate Hippo pathway regulation. Science 385, eadf4478. 10.1126/science.adf4478.

9. Lasker, K., Boeynaems, S., Lam, V., Scholl, D., Stainton, E., Briner, A., Jacquemyn, M., Daelemans, D., Deniz, A., Villa, E., et al. (2022). The material properties of a bacterial-derived biomolecular condensate tune biological function in natural and synthetic systems. Nat Commun 13, 5643. 10.1038/s41467-022-33221-z.

10. Su, X., Ditlev, J.A., Hui, E., Xing, W., Banjade, S., Okrut, J., King, D.S., Taunton, J., Rosen, M.K., and Vale, R.D. (2016). Phase separation of signaling molecules promotes T cell receptor signal transduction. Science 352, 595–599. 10.1126/science.aad9964.

11. Boyd-Shiwarski, C.R., Shiwarski, D.J., Griffiths, S.E., Beacham, R.T., Norrell, L., Morrison, D.E., Wang, J., Mann, J., Tennant, W., Anderson, E.N., et al. (2022). WNK kinases sense molecular crowding and rescue cell volume via phase separation. Cell 185, 4488–4506.e20. 10.1016/j.cell.2022.09.042.

12. Fujioka, Y., Alam, J.M., Noshiro, D., Mouri, K., Ando, T., Okada, Y., May, A.I., Knorr, R.L., Suzuki, K., Ohsumi, Y., et al. (2020). Phase separation organizes the site of autophagosome formation. Nature 578, 301–305. 10.1038/s41586-020-1977-6.

13. Wang, L., Choi, K., Su, T., Li, B., Wu, X., Zhang, R., Driskill, J.H., Li, H., Lei, H., Guo, P., et al. (2022). Multiphase coalescence mediates Hippo pathway activation. Cell 185, 4376–4393.e18. 10.1016/j.cell.2022.09.036.

14. Zhang, J.Z., Lu, T.-W., Stolerman, L.M., Tenner, B., Yang, J.R., Zhang, J.-F., Falcke, M., Rangamani, P., Taylor, S.S., Mehta, S., et al. (2020). Phase Separation of a PKA Regulatory Subunit Controls cAMP Compartmentation and Oncogenic Signaling. Cell 182, 1531–1544.e15. 10.1016/j.cell.2020.07.043.

15. Sampson, J., Richards, M.W., Choi, J., Fry, A.M., and Bayliss, R. (2021). Phase-separated foci of EML4-ALK facilitate signalling and depend upon an active kinase conformation. EMBO Rep 22, e53693. 10.15252/embr.202153693.

16. Tulpule, A., Guan, J., Neel, D.S., Allegakoen, H.R., Lin, Y.P., Brown, D., Chou, Y.-T., Heslin, A., Chatterjee, N., Perati, S., et al. (2021). Kinase-mediated RAS signaling via membraneless cytoplasmic protein granules. Cell 184, 2649–2664.e18. 10.1016/j.cell.2021.03.031.

17. Zhu, T., Xie, J., He, H., Li, H., Tang, X., Wang, S., Li, Z., Tian, Y., Li, L., Zhu, J., et al. (2023). Phase separation underlies signaling activation of oncogenic NTRK fusions. Proceedings of the National Academy of Sciences 120, e2219589120. 10.1073/pnas.2219589120.

18. Hardy, J.C., Pool, E.H., Bruystens, J.G.H., Zhou, X., Li, Q., Zhou, D.R., Palay, M., Tan, G., Chen, L., Choi, J.L.C., et al. (2024). Molecular determinants and signaling effects of PKA RIα phase separation. Molecular Cell 84, 1570–1584.e7. 10.1016/j.molcel.2024.03.002.

19. Park, J.-E., Zhang, L., Bang, J.K., Andresson, T., DiMaio, F., and Lee, K.S. (2019). Phase separation of Polo-like kinase 4 by autoactivation and clustering drives centriole biogenesis. Nat Commun 10, 4959. 10.1038/s41467-019-12619-2.

20. Sang, D., Shu, T., Pantoja, C.F., Ibáñez De Opakua, A., Zweckstetter, M., and Holt, L.J. (2022). Condensed-phase signaling can expand kinase specificity and respond to macromolecular crowding. Molecular Cell 82, 3693–3711.e10. 10.1016/j.molcel.2022.08.016.

21. Reinhardt, R., and Leonard, T.A. A critical evaluation of protein kinase regulation by activation loop autophosphorylation. eLife 12, e88210. 10.7554/eLife.88210.

22. Beenstock, J., Mooshayef, N., and Engelberg, D. (2016). How Do Protein Kinases Take a Selfie (Autophosphorylate)? Trends Biochem Sci 41, 938–953. 10.1016/j.tibs.2016.08.006.

23. Mitra, S.K., Hanson, D.A., and Schlaepfer, D.D. (2005). Focal adhesion kinase: in command and control of cell motility. Nat Rev Mol Cell Biol 6, 56–68. 10.1038/nrm1549.

24. Schaller, M.D., Hildebrand, J.D., and Parsons, J.T. (1999). Complex Formation with Focal Adhesion Kinase: A Mechanism to Regulate Activity and Subcellular Localization of Src Kinases. Mol Biol Cell 10, 3489–3505. 10.1091/mbc.10.10.3489.

25. Case, L.B., De Pasquale, M., Henry, L., and Rosen, M.K. (2022). Synergistic phase separation of two pathways promotes integrin clustering and nascent adhesion formation. eLife 11, e72588. 10.7554/eLife.72588.

26. Toutant, M., Costa, A., Studler, J.-M., Kadaré, G., Carnaud, M., and Girault, J.-A. (2002). Alternative splicing controls the mechanisms of FAK autophosphorylation. Mol Cell Biol 22, 7731–7743. 10.1128/MCB.22.22.7731-7743.2002.

27. Peeples, W., and Rosen, M.K. (2021). Mechanistic dissection of increased enzymatic rate in a phase-separated compartment. Nat Chem Biol 17, 693–702. 10.1038/s41589-021-00801-x.

28. Alberti, S., Gladfelter, A., and Mittag, T. (2019). Considerations and challenges in studying liquid-liquid phase separation and biomolecular condensates. Cell 176, 419–434. 10.1016/j.cell.2018.12.035.

29. Brami-Cherrier, K., Gervasi, N., Arsenieva, D., Walkiewicz, K., Boutterin, M.-C., Ortega, A., Leonard, P.G., Seantier, B., Gasmi, L., Bouceba, T., et al. (2014). FAK dimerization controls its kinase-dependent functions at focal adhesions. EMBO J 33, 356–370. 10.1002/embj.201386399.

30. Hsu, C.-P., Aretz, J., Hordeichyk, A., Fässler, R., and Bausch, A.R. (2023). Surface-induced phase separation of reconstituted nascent integrin clusters on lipid membranes. Proceedings of the National Academy of Sciences 120, e2301881120. 10.1073/pnas.2301881120.

31. Alanko, J., and Ivaska, J. (2016). Endosomes: Emerging Platforms for Integrin-Mediated FAK Signalling. Trends in Cell Biology 26, 391–398. 10.1016/j.tcb.2016.02.001.

32. Shen, Y., and Schaller, M.D. (1999). Focal Adhesion Targeting: The Critical Determinant of FAK Regulation and Substrate Phosphorylation. Mol Biol Cell 10, 2507–2518. 10.1091/mbc.10.8.2507.

33. van Mierlo, G., Jansen, J.R.G., Wang, J., Poser, I., van Heeringen, S.J., and Vermeulen, M. (2021). Predicting protein condensate formation using machine learning. Cell Reports 34, 108705. 10.1016/j.celrep.2021.108705.

34. Hou, S., Hu, J., Yu, Z., Li, D., Liu, C., and Zhang, Y. (2024). Machine learning predictor PSPire screens for phase-separating proteins lacking intrinsically disordered regions. Nat Commun 15, 2147. 10.1038/s41467-024-46445-y.

35. Chen, Z., Hou, C., Wang, L., Yu, C., Chen, T., Shen, B., Hou, Y., Li, P., and Li, T. (2022). Screening membraneless organelle participants with machine-learning models that integrate multimodal features. Proceedings of the National Academy of Sciences 24, e2115369119. 10.1073/pnas.2115369119.

36. Brasher, B.B., and Etten, R.A.V. (2000). c-Abl Has High Intrinsic Tyrosine Kinase Activity That Is Stimulated by Mutation of the Src Homology 3 Domain and by Autophosphorylation at Two Distinct Regulatory Tyrosines. Journal of Biological Chemistry 275, 35631–35637. 10.1074/jbc.M005401200.

37. Bonello, T.T., Cai, D., Fletcher, G.C., Wiengartner, K., Pengilly, V., Lange, K.S., Liu, Z., Lippincott-Schwartz, J., Kavran, J.M., and Thompson, B.J. (2023). Phase separation of Hippo signalling complexes. EMBO J 42, e112863. 10.15252/embj.2022112863.

38. Praskova, M., Khoklatchev, A., Ortiz-Vega, S., and Avruch, J. (2004). Regulation of the MST1 kinase by autophosphorylation, by the growth inhibitory proteins, RASSF1 and NORE1, and by Ras. Biochemical Journal 381, 453–462. 10.1042/BJ20040025.

39. Ambadi Thody, S., Clements, H.D., Baniasadi, H., Lyon, A.S., Sigman, M.S., and Rosen, M.K. (2024). Small-molecule properties define partitioning into biomolecular condensates. Nat. Chem. 16, 1794–1802. 10.1038/s41557-024-01630-w.

40. Kilgore, H.R., Mikhael, P.G., Overholt, K.J., Boija, A., Hannett, N.M., Van Dongen, C., Lee, T.I., Chang, Y.-T., Barzilay, R., and Young, R.A. (2024). Distinct chemical environments in biomolecular condensates. Nat Chem Biol 20, 291–301. 10.1038/s41589-023-01432-0.

41. Kota, D., Prasad, R., and Zhou, H.-X. (2024). Adenosine Triphosphate Mediates Phase Separation of Disordered Basic Proteins by Bridging Intermolecular Interaction Networks. J. Am. Chem. Soc. 146, 1326–1336. 10.1021/jacs.3c09134.

42. Bhatnagar, D., Roskoski, R.Jr., Rosendahl, M.S., and Leonard, N.J. (1983). Adenosine cyclic 3’,5’-monophosphate dependent protein kinase: a new fluorescence displacement titration technique for characterizing the nucleotide binding site on the catalytic subunit. Biochemistry 22, 6310–6317. 10.1021/bi00295a042.

43. Becher, I., Savitski, M.M., Savitski, M.F., Hopf, C., Bantscheff, M., and Drewes, G. (2013). Affinity Profiling of the Cellular Kinome for the Nucleotide Cofactors ATP, ADP, and GTP. ACS Chem. Biol. 8, 599–607. 10.1021/cb3005879.

44. Meyrat, A., and von Ballmoos, C. (2019). ATP synthesis at physiological nucleotide concentrations. Sci Rep 9, 3070. 10.1038/s41598-019-38564-0.

45. Patel, A., Malinovska, L., Saha, S., Wang, J., Alberti, S., Krishnan, Y., and Hyman, A.A. (2017). ATP as a biological hydrotrope. Science 356, 753–756. 10.1126/science.aaf6846.

46. Kim, T.H., Payliss, B.J., Nosella, M.L., Lee, I.T.W., Toyama, Y., Forman-Kay, J.D., and Kay, L.E. (2021). Interaction hot spots for phase separation revealed by NMR studies of a CAPRIN1 condensed phase. Proceedings of the National Academy of Sciences 118, e2104897118. 10.1073/pnas.2104897118.

47. Symons, M.H., and Mitchison, T.J. (1991). Control of actin polymerization in live and permeabilized fibroblasts. J Cell Biol 114, 503–513. 10.1083/jcb.114.3.503.

48. Lin, Y.-H., Kim, T.H., Das, S., Pal, T., Wessén, J., Rangadurai, A.K., Kay, L.E., Forman-Kay, J.D., and Chan, H.S. (2025). Electrostatics of salt-dependent reentrant phase behaviors highlights diverse roles of ATP in biomolecular condensates. eLife 13, RP100284. 10.7554/eLife.100284.

49. Aryan, F., Detrés, D., Luo, C.C., Kim, S.X., Shah, A.N., Bartusel, M., Flynn, R.A., and Calo, E. (2023). Nucleolus activity-dependent recruitment and biomolecular condensation by pH sensing. Molecular Cell 83, 4413–4423.e10. 10.1016/j.molcel.2023.10.031.

50. J. Kathiriya, J., Ramesh Pathak, R., Clayman, E., Xue, B., N. Uversky, V., and Davé, V. (2014). Presence and utility of intrinsically disordered regions in kinases. Molecular BioSystems 10, 2876–2888. 10.1039/C4MB00224E.

51. Ou, X., Lao, Y., Xu, J., Wutthinitikornkit, Y., Shi, R., Chen, X., and Li, J. (2021). ATP Can Efficiently Stabilize Protein through a Unique Mechanism. JACS Au 1, 1766–1777. 10.1021/jacsau.1c00316.

52. Chuang, H.-H., Zhen, Y.-Y., Tsai, Y.-C., Chuang, C.-H., Hsiao, M., Huang, M.-S., and Yang, C.-J. (2022). FAK in Cancer: From Mechanisms to Therapeutic Strategies. Int J Mol Sci 23, 1726. 10.3390/ijms23031726.

53. Wang, J., and Pendergast, A.M. (2015). The Emerging Role of ABL Kinases in Solid Tumors. Trends Cancer 1, 110–123. 10.1016/j.trecan.2015.07.004.

54. Mitrea, D. M., Mittasch, M., Gomes, B. F., Klein, I. A., and Murcko, M. A. (2022). Modulating biomolecular condensates: a novel approach to drug discovery. Nature Reviews Drug Discovery 21, 841–862.

55. Potter, M., Newport, E., and Morten, K.J. (2016). The Warburg effect: 80 years on. Biochem Soc Trans 44, 1499–1505. 10.1042/BST20160094.

