## Supplemental Material for "Kinase condensates enrich ATP and trigger autophosphorylation"

**The Supplemental file includes:**

Materials and Methods
Supplementary Text
Figs. S1 to S15
Table S1
Supplemental References

**Other Supplemental Materials for this manuscript include the following:**

Data S1

### Materials and Methods

Microscopy

TIRF images were captured using LASX acquisition software to drive a Leica TIRF-module mounted on a Leica DMi8 with DIC optics equipped with a plan apo 100 Å~ 1.47 NA TIRF objective and a Quad Band set filter cube for TIRF applications. Illumination was provided by an integrated laser system equipped with multiple laser lines (405 nm-50mw/488 nm-150mw/561 nm-120mw/638 nm-150mW). Images were acquired using a Hamamatsu Flash 4.0 V3 CMOS camera.

Epifluorescence and DIC images were captured using LASX acquisition software to drive a Leica DMi8 with DIC optics equipped with a plan apo 100 Å~ 1.47 NA TIRF objective. Illumination was provided by an LED8 light source equipped with two interchangeable filter cubes (one for excitation at 391/32, 479/33, 554/24, 638/31; one for excitation at 473/22, 539/24, 641/78, 810/80). Images were acquired using a Hamamatsu Flash 4.0 V3 CMOS camera.

Spinning disk confocal images were captured using MetaMorph acquisition software to drive a Zeiss AxioVert 200M inverted microscope stand with DIC optics equipped with a Yokogawa CSU-22 spinning disk confocal scan head with Andor Borealis modification. Illumination was provided by an Andor integrated laser engine equipped with multiple laser lines (405nm, 445nm, 488nm, 515nm, 561nm, and 642nm). Images were acquired using a Hamamatsu Orca-ER cooled CCD camera.

Preparing PEG-Silane coated plates

384-well glass-bottom plate (Cellvis, P384-1.5H-N) was washed with 5% Hellmanex III (Hëlma Analytics) for 3.5 hrs at 55 °C and thoroughly rinsed with MilliQ $H_2O$. Plates were then washed with 1 M NaOH for 1 hr at 55 °C and thoroughly rinsed with MilliQ $H_2O$. 50 µ L of 20 mg/mL mPEG silane MW 5 k (Creative PEGworks) in 95% EtOH was added to each well. The plate was covered in parafilm and incubated overnight at room temperature. The plate was thoroughly rinsed with MilliQ $H_2O$, dried, and sealed with foil.

Microscopy of condensates

Representative images of in vitro condensates were taken with epifluorescence or differential interference contrast (DIC) microscopy. The buffer conditions used for specific experiments are denoted in figure legends. For all experiments, stock proteins were thawed quickly at room temperature then spun at 21,000 rpm at 4$^{o}$C for 10 minutes to remove any aggregates. The supernatant was transferred to a new tube and A280 values were measured using the protein storage buffer as a blank. Protein concentrations were calculated based on predicted extinction coefficients. If necessary, protein stocks were diluted in their respective storage buffer before being diluted into an experimental buffer that yielded the desired final concentrations of all buffer components (buffer, salt, etc.) after accounting for contributions from the protein storage buffers. For all experiments, protein was always added after buffer with the protein/s required for phase separation added last. The solution was mixed by pipetting up and down before loading solutions into a 384-well PEG-silane coated glass-bottom plate. Samples were incubated for 20-

30 minutes before imaging. For each experiment, regions of interest were randomly selected for imaging, and all images were acquired within 10 minutes of each other. Fluorescence microscopy images were analyzed in ImageJ[1] to calculate the percent droplet area. The background was subtracted with a rolling circle of radius 25 pixels, then hot/dead pixels were removed by using a median filter with pixel size=2. Then any fields of view (FOV) with large auto-fluorescent debris were removed. Images were auto-thresholded with the Li algorithm and "Analyze Particles" was run to calculate percent area covered by droplets for each image. For DIC microscopy analysis droplets were thresholded by training a Pixel Classification segmentation algorithm in Illastik on one image to be analyzed then used to segment the remaining images. "Analyze Particles" was run in ImageJ to calculate percent area covered by droplets for each image. For all microscopy experiments, representative images for figures were selected such that the percent area of the representative image closely matched the average of all the images for that condition.

Turbidity measurements

50 μL phase separated solutions were prepared as described for the microcopy experiments and incubated for 45 minutes. After incubation, samples were gently mixed by pipetting 3-4 times with a 200 μL pipette tip and then carefully transferred into a low volume quartz cuvette. Absorbance at 350 nM was measured, and the final buffer without protein was used as the blank measurement.

FAK phase separation autophosphorylation assays

FAK proteins were incubated in 50 mM HEPES pH 7.5, 50 or 300 mM NaCl (low or high salt conditions), 250 μM $MnCl_2$, 0.435% (v/v) glycerol, and 1 mM DTT. A 100 μL mixture was made and 15 μL of this mixture was quickly aliquoted into 7 PCR tubes, one for each timepoint. Mixtures were incubated for 45 minutes at room temperature to reach equilibrium. 0.75 μL of a 100 mM stock of ATP in 100 mM Tris-HCl pH 7.5 was added to 999.25 μL of 25 mM HEPES pH 7.5, 50mM NaCl, 1 mM DTT to yield a 75 μM ATP solution. To initiate the reactions, 2 μL of the 75 μM ATP solution was added to the FAK mixture and instantly mixed 3-4 times by pipetting up and down with a 10 μL pipette tip. The reactions were quenched by adding 4.5 μL of Quench Buffer (150 g/L SDS, 0.3 M Tris pH 6.8, 25% (v/v) glycerol) preheated to 99 $^{o}$C and instantly mixed 3-4 times by pipetting up and down with a 10 μL pipette tip. Reactions were then incubated at 99 $^{o}$C for 5 minutes, then placed on ice. Reactions were spun down in a tabletop centrifuge to collect all the samples to the bottom of the tube and 5 μL was diluted with 95 μL of 2X SDS Loading Buffer (100 mM Tris pH 6.8, 10% (v/v) 2-mercapto-ethanol, 4% (w/v) SDS, 0.2% (w/v) Bromophenol Blue, 20% (v/v) glycerol). This dilution was determined to load samples in the middle of the linear range of both the pan-FAK and pY397 FAK channels for the western blot protocol. Samples were analyzed by western blot. Primary antibodies were mouse Anti-FAK, clone 4.47 (Sigma; 1:2000) and rabbit Anti-Phospho-FAK (Tyr397) (Invitrogen; 1:2000). Secondary antibodies were goat Anti-mouse IgG (H+L) (DyLight™ 680 Conjugate; 1:10,000) (Cell Signaling Technology, #5470) and goat Anti-rabbit IgG (H+L) (DyLight™ 800 4X PEG Conjugate: 1:10,000) (Cell Signaling Technology, #5151).

All blots were analyzed in ImageJ using the same region of interest (ROI) size and shape. This ROI was moved to surround each band in the pan-FAK channel and pasted onto the same position in the pY397 channel to measure the average intensity of the same band in both channels. Background measurements were taken from areas just below or above each band. For each band, its associated background measurement was subtracted from the average intensity. Finally, the pY397 value was divided by the pan-FAK value for each timepoint to generate the normalized pY397 signal in figure 1 C.

<u>FAK total vs dilute autophosphorylation assays</u>

Assays were performed as described above in “FAK Phase Separation Autophosphorylation Assays” with the following modifications. A 200 μL solution of 1 μM dephosphorylated FAK in a final buffer of 50 mM HEPES pH 7.5, 50 mM NaCl, 250 μM MnCl2, 1% glycerol, 1 mM DTT was made in a 1 mL ultracentrifuge tube (Beckman, #347356). This was immediately mixed, and half was removed to a separate ultracentrifuge tube. These were incubated or 1 hour at room temperature to reach equilibrium. One tube (dilute sample) was then spun down at 40,000 rpm at 22°C for 3 hours to remove the dense phase. 50 μL of the supernatant was removed as dilute phase. The tube that was not centrifuged was incubated stationary at room temperature then mixed by pipetting up and down gently 4-5 times to resuspend any settled droplets. Reactions were run as described above for 30 second timepoints and 0 second timepoints with no ATP added. Quenched samples were diluted appropriately with 2X SDS Loading Buffer (100 mM Tris pH 6.8, 10% (v/v) 2-mercapto-ethanol, 4% (w/v) SDS, 0.2% (w/v) Bromophenol Blue, 20% (v/v) glycerol) to a final concentration of ~7 nM FAK (3.025-fold dilution of dilute samples and 100-fold dilution of total samples). Samples were run along with 6 phosphorylated FAK standards. These standards were made from purified phosphorylated FAK diluted in protein storage buffer. The final concentrations of the standards were (before dilution in 2X SDS Loading Buffer) 0.977 nM, 1.953 nM, 3.906 nM, 7.8125 nM, 15.625 nM, and 31.25 nM. Samples were analyzed by western blot using the antibodies described above.

For all bands measurement and background subtraction was performed as described in “FAK Phase Separation Autophosphorylation Assays”. The FAK and pFAK measurements of the standards were plotted as amount loaded in femtomoles vs average intensity. These plots were fit with a linear regression, and the equation of this line was used to generate absolute amounts of FAK and pY397 for each sample. The pY397 FAK amount was normalized by dividing by the FAK amount for each sample. The resulting value of the 0 second timepoint was subtracted from the 30 second timepoint and this was divided by 30 seconds to yield the rate. The values were then multiplied by their respective dilution factors (100 for total samples and 3.025 for dilute samples). These values were then multiplied by 100 to generate arbitrary units for ease of displaying the data.

<u>FAK titration autophosphorylation assays</u>

Assays were performed as above in “FAK Phase Separation Autophosphorylation Assays” with the following modifications. 25 μL solutions of dephosphorylated GFP-FAK at indicated concentrations (10-80 nM) were made in a final buffer of 25 mM HEPES pH 7.5, 50 mM NaCl, 1% glycerol, 1 mM DTT in Low Protein Binding Microcentrifuge Tubes (Thermo) and incubated for 2 hours at room temperature. 2.5 μL of a 25 mM HEPES pH 7.5, 50 mM NaCl, 1 mM DTT, 1.1 mM ATP, 2.75 mM $MnCl_2$ solution was added to initiate reactions. ATP wasn't added to one sample to indicate starting phosphorylation amount. Reactions were incubated at room temperature for 30 seconds. Reactions were quenched with 6.88 μL of Quench Buffer. Quenched samples were diluted appropriately with 2X SDS Loading Buffer (100 mM Tris pH 6.8, 10% (v/v) 2-mercapto-ethanol, 4% (w/v) SDS, 0.2% (w/v) Bromophenol Blue, 20% (v/v) glycerol) to a final concentration of 3.636 nM mEGFP-FAK. Samples were run along with 8 phosphorylated FAK standards. These standards were made from purified phosphorylated FAK diluted in protein storage buffer. The final concentrations of the standards were (before dilution in 2X SDS Loading Buffer) 0.244 nM, 0.488 nM, 0.977 nM, 1.953 nM, 3.906 nM, 7.8125 nM, 15.625 nM, and 31.25 nM.

For all bands, measurement, background subtraction, standard curve generation, and pY397 normalization to total FAK was performed as described in "FAK Total vs Dilute Autophosphorylation Assays" above. These values were then multiplied by their respective dilution factors to yield the plotted values.

Western blotting

10 μL of each sample was loaded on a 10% Tris/Glycine SDS-PAGE gel and run at 240V for 30 minutes. 5 μL of a 1:100 dilution of Color Prestained Protein Standard, Broad Range (10-250 kDa) (NEB) in 2X SDS Loading Buffer was loaded as well. Additionally, for autophosphorylations assays a control sample with 2 μL of water added in lieu of the ATP solution was run to show the phosphorylation state of the starting protein. SDS-PAGE gels were washed quickly in TBS then assembled into transfer sandwich with a Low Fluorescence PVDF membrane pre-wet in 100% methanol and 5 pieces of Whatman paper (per side) pre-soaked in 1X Trans-Blot Transfer buffer (BIO-RAD). Transfer was run in a Trans-Blot Turbo (BIO-RAD) with MIXED setting. Membranes were then washed quickly in TBS then transferred to an incubation box with 5 mL of Azure Fluorescent Blot Blocking Buffer (Azure Biosystems). Membranes were incubated at room temperature with gentle shaking for 1 hour to block. Primary antibodies were added at indicated dilutions and incubated at $4^{o}$C overnight with gentle shaking. Membranes were washed 3 times with 20 mL Azure Fluorescent Blot Washing Buffer (Azure Biosystems). Membranes were incubated with 5 mL of secondary antibodies in Azure Fluorescent Blot Blocking Buffer at room temperature with gentle agitation for 1 hour. The membrane was transferred to a new incubation box with 20 mL of Azure Fluorescent Blot Washing Buffer and quickly washed. This step was repeated once, then the membrane was transferred to a new incubation box with 20 mL of Azure Fluorescent Blot Washing Buffer and incubated at RT with gentle shaking for 5 minutes. The wash buffer was decanted, and the blot was washed again for 5 mins. This was repeated for 3 total 5-minute washes. The membrane was transferred to a new incubation box with 20 mL of TBS and quickly washed. The membrane was transferred to a new incubation box with 20 mL of TBS and then imaged using a ChemiDoc MP Imaging System (BIO-RAD).

FAK dilute and dense phase concentration measurements via microscopy

Concentrations of mEGFP-FAK in both the dilute and dense phases were measured by using a standard curve of dephosphorylated mEGFP-FAK in a high salt buffer that prevents phase separation. These standards were 200 nM, 100 nM, 50 nM, 25 nM and 12.5 nM in High Salt Buffer (25 mM HEPES pH 7.5, 500 mM NaCl, 1 mM DTT, 20 mM glucose, 800 µg/mL glucose oxidase (Sigma, G2133), 140 µg/mL catalase (Sigma, C1345)). For samples to measure the dense phase, 0.1% dephosphorylated GFP-FAK was used at 2 µM in Low Salt Buffer (25 mM HEPES pH 7.5, 50 mM NaCl, 1 mM DTT, 20 mM glucose, 800 µg/mL glucose oxidase (Sigma, G2133), 140 µg/mL catalase (Sigma, C1345)). This 0.1% GFP-FAK was created by mixing the appropriate amount of GFP-FAK and untagged FAK stocks. For samples to measure the dilute phase, 100% dephosphorylated GFP-FAK was used at 50 nM in Low Salt Buffer. These samples, along with the blanks of Low Salt Buffer and High Salt Buffer, were loaded and imaged in wells in a 384-well glass bottom plate that were prepared as described below.

Wells were previously cleaned and coated with PEG-Silane and then incubated with 5 µL of a 1:20,000 dilution of 0.5 nm red FluoSpheres™ (Invitrogen™) in 100% ethanol for 10 minutes at room temperature until dry. Wells were then incubated with 80 µL of Blocking Buffer (25 mM HEPES pH 7.5, 50 mM NaCl, 1 mM DTT, 1mg/mL BSA) for 30 minutes at room temperature, then washed quickly 3 times with 100 µL of either High Salt Buffer or Low Salt Buffer immediately preceding sample loading. Samples were loaded then incubated for 30-90 minutes while setting up the microscope and imaging of all wells was completed within 2 hours. Wells were imaged with a spinning disc confocal microscope. Red fluorospheres were used to focus the microscope for all measurements.

Standard curves for determining dense and dilute phase concentrations were generated as follows. For each of the standards and the High Salt Buffer blank well, ~ 10 images of each well were averaged together to create an average image. Any images that showed debris or unusual fluorescence were omitted. The average images of the standards were corrected for uneven illumination by dividing by the average image of the High Salt Buffer sample and then multiplying by the mean pixel intensity of the average image of the High Salt Buffer. Then the mean intensity of each corrected standard was measured, and the mean intensity of the High Salt Buffer (background) was subtracted. These values were fit to a linear regression to create a standard curve. New standards were imaged with each set of experiments.

Dilute phase sample images were corrected for uneven illumination as described above. Images were then thresholded manually to select the regions excluded by the droplets, and the average intensity of this region was measured. The Low Salt Buffer sample was averaged, corrected for uneven illumination, and measured as described above. This background value was subtracted from all of the dilute phase measurements. The resulting background subtracted measurements were converted to dilute phase concentrations using the linear regression of the standard curve.

For dense phase measurements, Z-stacks with width 0.5 µm were taken. The dense phase images were analyzed similarly with the following exceptions. Uneven illumination and background subtraction were performed using the Low Salt Buffer blank. Z-series of each field of view were thresholded for individual 3D droplets using the 3D Object Counter plugin. Threshold settings were determined independently for each replicate. The mean intensity of each 3D droplet was taken to calculate the concentration. Since point spread function and photobleaching affects would lower mean intensity values in the droplets, this is a conservative lower bound estimate of the droplet concentration. Background from the Low Salt Buffer blank was subtracted from these mean intensity values and then the dense phase concentration was calculated from the linear regression of the standard curve. Since these experiments used only 0.1% GFP-FAK (while the standards were made from 100% GFP-FAK) the resulting concentrations were multiplied by 1000 to yield the true dense phase concentration.

Mass photometry

Samples of protein were diluted as specified in the figure legends and incubated at room temperature for 2 hours in Low Protein Binding Tubes (Thermo). Buffers were filtered twice using 0.22 μm syringe filters before use. Standards were prepared by diluting NativeMark™ Unstained Protein Standard (Invitrogen) 100-fold in identical buffer to samples. Samples and standards were run on the Mass Photometer (Refeyn) using buffer-less focusing. Once samples were loaded, samples were checked for proper focus and adjusted manually and moved to a new region if needed. This procedure only analyzes the dilute phase, since fields of view containing condensates would fail the focus procedure. Contrast values for standards were matched to known molecular weights, and this was used to compute mass values of samples in the Refeyn software.

Cell culture and transfection

Mouse embryonic fibroblasts (MEFs, ATCC CRL-2645) were passaged according to the ATCC recommendations. MEFs were cultured in Dulbecco's Modified Eagle Medium (DMEM) supplemented with 10% Fetal Bovine Serum (FBS), Penicillin/Streptomycin (100 U/mL and 100 μg/mL, respectively) and Glutamax (2 mM) at 37°C with 5% $CO_2$. Cells were routinely tested for mycoplasm. All transient transfections were performed with Invitrogen™ Lipofectamine™ 3000 Transfection Reagent following the manufacturer's protocol. MEFs were transfected with 8 μg of plasmid DNA at 50% confluency in a 6-well plate. After 24 hours, cells were trypsinized and 50,000 cells were seeded into 8-well iBidi chambers coated with poly-D-Lysine. Cells were imaged by epifluorescence and differential interference contrast (DIC) microscopy at 37°C with 5% $CO_2$. Poly-D-Lysine coated plates were prepared with poly-D-Lysine (Thermo Scientific A3890401) following the manufacturer's protocol. Fibronectin coated plates were made by diluting Bovine Fibronectin (Sigma Aldrich #F1141) 100-fold in Dulbecco's Phosphate Buffered Saline (DPBS) and incubating wells at 4°C overnight. This solution was removed, and wells were washed once in DPBS immediately before use.

Generation of stable DOX-inducible GFP-FAK-WT cell line

MEFs were co-transfected with XLone-Puro-mEGFP-FAK-WT (cargo) and Super PiggyBac Transposase plasmid (Gift of Calo lab) in 6-well plates[2]. After 3 days (to allow insertion of the cargo and to allow loss of unincorporated cargo plasmid), cells were expanded to 10 cm plates and cells with successful incorporation and expression of the cargo were selected with 10 μg/mL Puromycin. Control/untransfected cells were also selected for in parallel to assess when selection was successful by noting when all control cells died. After this period (roughly 3-5 days), cells were expanded to 40 million cells and Flow Assisted Cell Sorted (FACS) sorted. The top 1% GFP expressing cells were collected and expanded to 40 million cells. At this point cells were tested for mycoplasma and aliquots were frozen for storage. For all experiments, mEGFP-FAK-WT was induced with 4 μg/μL of Doxycycline hyclate (Sigma-Aldrich, #D5207) for 24 hours.

Fluorescence recovery after photobleaching (FRAP) and FRAP analysis

Puncta/condensates were bleached with exposure to 488 nm laser at 80% power recovery was imaged at timepoints indicated in graphs. FRAP images were analyzed using a region of interest (ROI) outside of the cell as the background signal, an ROI surrounding the entire cell as the total signal, and an ROI surrounding the mEGFP-FAK puncta as the puncta signal. The background signal was subtracted from both the puncta and total signal, and the resulting background-subtracted puncta signal was divided by the background-subtracted total signal to correct for photobleaching and laser intensity fluctuations over time. These values were then normalized so the initial (pre-bleach) signal was 1 in all replicates. For each replicate, the intensity of the first timepoint coinciding with the bleach event ($I_B$) was subtracted from all time points, and the resulting data was fit to a single exponential regression to determine $t_{1/2}$. Percent recovery was calculated by dividing the plateau value extracted from the regression by ($1-I_B$). FRAP analysis of mEGFP-FAK condensates *in vitro* was performed similarly, except using an unbleached droplet as the total ROI to control for photobleaching and laser intensity fluctuations over time.

<u>Puncta vs cell intensity analysis</u>

Cells transfected with indicated constructs were plated on poly-D-Lysine as described. 15x15 field of view (FOV, 132x132 micron each) tilescans were imaged with DIC microscopy and epifluorescence microscopy, acquiring Z-stacks of each FOV to capture the entire volume of cells. Maximum intensity projections of each FOV were created, and the FOVs were stitched using LASX software. Individual cells were cropped in ImageJ and autothresholded with Otsu algorithm. Mean fluorescence intensity of each cell was measured using ImageJ "Analyze Particles". The number of puncta in each cell were counted manually. Cells with large amorphous aggregates were excluded from the puncta analysis (Data on aggregates can be found in Supplemental Figure S5). The total number of cells in the tilescan (both transfected and untransfected) was estimated by counting cells in the DIC channel, and percent of total cells with puncta was calculated for western blot interpretation. For conditions co-expressing mCherry-Paxillin analysis was limited to cells expressing a small range of mCherry signal (300-600 raw intensity values) which was empirically determined to be the smallest range that still captured a wide range of GFP signals. This keeps the paxillin concentration relatively constant in the analyzed cells, enabling measurement of the specific effect of increasing mEGFP-FAK concentrations.

<u>Cell lysate western blots</u>

MEFs were either transfected with 8 μg of plasmid DNA, mock transfected (No DNA), or untreated at 70-80% confluency in a 6-well plate. After 24 hours, all cells were trypsinized and seeded into 6-well plates coated with Poly-D-Lysine for 30 minutes. One sample (ECM) of untreated cells was not trypsinized or re-plated to act as a reference for normal integrin-activated pY397 FAK/pan-FAK (or pCas/Cas or pPax/Pax) ratios. At this point, cells were washed twice in ice-cold phosphate buffered saline (PBS) and lysed with 100 μL of ice-cold RIPA buffer (Thermo Scientific #89900) supplemented with 2.5mM sodium orthovanadate. Lysates were collected and stored at -20°C until used for western blots. Total protein concentration was measured by diluting a small amount of lysate in 25 mM HEPES pH 7.5 and using a Micro BCA assay (Thermo Scientific #23231) with albumin standards prepared in 25 mM HEPES pH 7.5. Dilutions of the reference sample were run on each western as both adhesion positive pFAK/FAK reference samples and on-blot standards to confirm linearity of pan-FAK and pY397 FAK signals. Samples were loaded accordingly to attain pan-FAK and pFAK signals in this linear range. Samples were diluted 2-fold in 2X SDS Loading Buffer (100 mM Tris pH 6.8, 10% 2-mercapto-ethanol, 4% SDS, 0.2% Bromophenol Blue, 20% glycerol) and boiled at 99°C for 10 minutes, then placed on ice for 5 minutes. 10 μL of each sample was run on a 10% Tris-Glycine gel for 35 minutes at 240V. Transfer, blocking and antibody incubation was performed identically as described earlier for FAK and pY397 FAK. For each blot a linear regression of the standards was used to determine the relative amount of pan-FAK and pY397 FAK in each sample. The pY397 FAK signal was divided by the pan-FAK signal for each sample and then normalized so the values of the reference samples equaled 1. These normalized values represent the percent of reference pY397/FAK ratios recovered by each condition. Also, because of the different mobilities of the GFP tagged FAK and endogenous FAK, it was possible to analyze both species separately on the same blot. For samples that were stained for total protein, this was done with the AzureRed Protein Stain (AC2124) from Azure Biosystems following the manufacturer's protocol. Lysates were also blotted for pCas/Cas and pPax/Pax analysis following the same protocol except using the following primary antibodies and dilutions; 1:1000 Mouse anti p130Cas (BD Transduction Laboratories™, Purified Mouse Anti-p130 [Cas] #610271), 1:1000 Rabbit anti pY410 p130 Cas (Cell Signaling Technology, Phospho-p130 Cas (Tyr410) Antibody #4011), 1:10,000 Mouse anti paxillin (BD Transduction Laboratories™, Purified Mouse Anti-Paxillin #612405), 1:1000 Rabbit anti pY118 paxillin (Cell Signaling Technology, Phospho-Paxillin (Tyr118) Antibody #2541).

Immunostaining

All immunostaining was performed using the protocol below. The following antibodies and concentrations were used. 1:100 Rabbit anti pY397 FAK (Invitrogen, Phospho-FAK (Tyr397) Polyclonal Antibody #44624G), 1:100 Rabbit anti EEA1 (Cell Signaling Technology, EEA1 Antibody #2411), 1:50 Rabbit anti Rab11 (Cell Signaling Technology, Rab11 (D4F5) XP® Rabbit mAb #5589), 1:250 AF647 Donkey anti Rabbit (Invitrogen, Donkey anti-Rabbit IgG (H+L) Highly Cross-Adsorbed Secondary Antibody, Alexa Fluor™ Plus 647, # A32795TR). Exogenously expressed mEGFP-FAK was imaged using the intrinsic GFP fluorescence.

All steps were carried out in 8-well iBidi microplates coated with Poly-D-Lysine. Because cell adhesion under these conditions is weak, care was taken to add and remove buffers slowly on the side of the wells to minimize cell loss. All steps performed at room temperature unless otherwise noted. 50,000 cells were seeded in wells for 10 minutes at 37°C and 5% $CO_2$ to allow adhesion to Poly-D-Lysine surface. Cells were then washed 2 times in PBS and fixed with 4% paraformaldehyde (PFA) in tris-buffered saline (TBS) for 15 minutes at 37°C. Cells were then permeabilized with 0.5% (v/v) TritonX-100 in TBS for 8 minutes without shaking. Cells were then washed with 0.1M Glycine in TBS for 10 minutes without shaking. Cells were then washed two times, 5 minutes then 10 minutes, with TBS supplemented with 0.1% (v/v) Tween 20 (TBS-T) with gentle shaking. Cells were then blocked with 2% (w/v) bovine serum albumin (BSA) in TBS-T for 1 hour with gentle shaking. Primary antibodies in 2% BSA in TBS-T were added and cells were incubated for 2 hours with gentle shaking. Cells were then washed three times with TBS-T for 5 minutes. Secondary antibody in 2% BSA in TBS-T were added and cells were incubated for 1 hour with gentle shaking. Cells were then washed three times with TBS-T for 5 minutes. Cells were then quickly washed one time with TBS, the TBS was replaced, and samples were imaged.

Mst2 and Abl western blots

To confirm kinase activity in standard kinase buffers, 5 μg of kinase was added to 1X NEBuffer™ for Protein Kinases (50 mM Tris-HCl, 10 mM $MgCl_2$, 0.1 mM EGTA, 2 mM DTT, 0.01% Brij 35 (pH 7.5 @ 25°C) and 25 μL reactions were initiated with 0.5 mM ATP. Reactions were incubated at room temperature for 2 hours. Mock reactions without ATP added were also set up to assess initial phosphorylation states. Samples were analyzed by western blotting. For MST2 the following primary antibodies were used: 1:1000 rabbit anti Phospho-MST1 (Thr183)/MST2 (Thr180) (Cell Signaling Technology, #49332) and 1:1000 mouse anti STK3 Monoclonal Antibody (4F7) (Abnova). For Abl1 the following antibodies were used: 1:1000 rabbit anti Phospho-c-Abl (Tyr245) Antibody (Cell Signaling Technology, #2861) and 1:1000 mouse anti c-Abl Antibody (24-11): sc-23 (Santa Cruz Biotechnology). Secondary antibodies were goat Anti-mouse IgG (H+L) (DyLight™ 680 Conjugate; 1:10,000) (Cell Signaling Technology, #5470) and goat Anti-rabbit IgG (H+L) (DyLight™ 800 4X PEG Conjugate: 1:10,000) (Cell Signaling Technology, #5151).

ADP-Glo™ kinase assays and analysis

Autophosphorylation assays were performed using purified kinases and the ADP-Glo™ Kinase Assay kit (Promega #V6930). Purified kinases were incubated at 1 μM final concentration in buffers to form condensates as described previously except buffer was supplemented with 0.5 mM $MgCl_2$. Samples were either centrifuged at 40,000 rpm at 22°C for 1 hour or incubated at 22°C for 1 hour. Supernatant was removed from centrifuged samples to serve as the dilute phase samples. The non-centrifuged samples were used as the total samples. The total samples were mixed by pipetting up and down to resuspend any settled condensates before initiating reactions. Reactions were initiated by adding ATP at a final concentration of 1 μM and immediately mixed. 5 μL samples were removed before adding ATP (0 second timepoint) and every 30 seconds for 2 minutes and mixed with 5 μL of ADP-Glo™ Reagent to stop the reaction and deplete unconsumed ATP. These solutions were incubated at RT for 40 minutes. Then, 10 μL of Kinase Detection Reagent was added and incubated at RT for 30 minutes. Luminescence of samples and a 1 μM series of ATP+ADP standards were measured using a BioTek Synergy H1 hybrid reader at room temperature. These ATP+ADP standards were generated according to the manufacturer's protocol using separate buffers that match each kinases phase separation buffer composition. In short, these standards represent different percentages of ATP conversion to ADP. The kit can only reliably detect down to 1-2% conversion of the total ATP to ADP. The percent of ATP converted to ADP was determined for each sample by using the equation of the linear regression of the ATP+ADP standards. This percentage was then converted to fmol ATP generated. For all samples, the signal of the 0 second timepoint was subtracted from each timepoint and the amount of ADP generated was divided by the seconds of each timepoint. For each sample, the timepoint that led to the largest rate was chosen to represent the intial rate. To be considered to have a detectable signal/rate above the noise the timecourse needed to have at least 3 consecutive points with increasing amounts of ADP generated and the largest signal of those 3 points represent a non-negative amount of ADP generated. If this was not achieved the samples were assumed to be below the limit of detection of the assay.

Phospho- mass spectrometry

Samples were digested on S-trap micro spin columns from Protifi. Samples were digested following the manufacturers protocol with the following changes:

Step 5: 10 mM DTT (final concentration) was used instead of TCEP to reduce the proteins and the samples were placed on a heating block for 10 minutes at 95C.

Step 6: 20 mM iodoacetamide (final concentration) was used instead of MMTS, and samples incubate at RT for 30 minutes in the dark.

Step 7: 1% Trifluoroacetic acid (final concentration) was used instead of phosphoric acid.

Step 14: Two separate enzymes were used Trypsin and Chymotrypsin and samples were digested overnight at 37C.

After digestion samples were desalted using Pierce Peptide Desalting Spin Columns (cat#89852) per manufacturer protocol. The tryptic peptides were loaded on a precolumn (Acclaim PepMap 100 75 μM x 2 cm) and separated by reverse phase HPLC (Thermo Ultimate 3000) using a Thermo PepMap RSLC C18 column (2 μm tip, 75 μm x 50 cm PN# ES903) over a gradient (below) before nano-electrospray using a Orbitrap Exploris 480 mass spectrometer (Thermo). Solvent A was 0.1% formic acid in water and solvent B was 0.1% formic acid in acetonitrile.

| Time | % B buffer | Flow rate nL |
|---|---|---|
| 0 | 1 | 200 |
| 15 | 1 | 200 |
| 15.5 | 3 | 200 |
| 35 | 23 | 200 |
| 41 | 35 | 200 |
| 43 | 80 | 200 |
| 46 | 80 | 200 |
| 46.1 | 1 | 200 |
| 60 | 1 | 200 |

The mass spectrometer was operated in a data-dependent mode. The parameters for the full scan MS were: resolution of 60,000 across 375-1500 *m/z* and maximum IT 25 ms. The full MS scan was followed by MS/MS for the top 15 precursor ions in each cycle with a NCE of 28, dynamic exclusion of 10 s and resolution of 30,000. Raw mass spectral data files were searched using Sequest HT in Proteome Discoverer (Thermo). Sequest search parameters were: 10 ppm mass tolerance for precursor ions; 0.02 Da for fragment ion mass tolerance; 2 missed cleavages of trypsin; fixed modification were carbamidomethylation of cysteine and peptide N-termini; variable modifications were methionine oxidation, tyrosine, serine and threonine phosphorylation, methionine loss at the N-terminus of the protein, acetylation of the N-terminus of the protein and also Met-loss plus acetylation of the protein N-terminus. Data was searched against the protein sequences of the kinases used. The canonical autophosphorylation site of Mst2 was not detected due to lack of coverage of that region with chymotrypsin or trypsin digestion.

Luciferase assays for measuring nucleotide concentration in dense and dilute phase

250 μL samples of phase separated proteins at concentrations and buffers indicated in figure legends were prepared in 1 mL ultracentrifuge tube (Beckman, #347356). Samples were mixed and incubated for 1 hour at room temperature to reach equilibrium. At this point nucleotides (ATP, ADP or GTP) were added at indicated final concentrations and mixed. Samples of poly-L-Lysine condensates were prepared similarly except nucleotides were added initially to induce phase separation. Samples were then spun down at 40,000 rpm at 22$^oC$ for 1 hour to pellet the dense phase. The supernatant was carefully removed and used for dilute phase samples. The dense phase was either collected by repeatedly tapping a pipette tip to force the viscous dense phase into the tip or, when too small to be manipulated this way, all supernatant was removed and dense phase sample was retained in the tube. For dense phase samples that were too small to be manipulated the volume was overestimated to be 1 μL for downstream calculations (which would underestimate concentration calculations later). For samples that could be packed into a pipette tip, the volume was calculated by using a 1 μL mark on the pipette tip as a reference and assuming the pipette tip was a cylinder. Dense phase samples were added to 10 μL of 6M guanidine HCl and mixed to dissolve. Dilute phase samples were diluted 1:10 in 6M guanidine HCl. 5 μL of these samples were then diluted a further 1:100 in 25 mM HEPES pH 7.5, 200 mM NaCl, 10 mM $MgCl_2$. Also, experiments with millimolar concentrations of nucleotide required one additional 1:100 (or 1:1000 for poly-L-Lysine dense phase) dilution step in 6M guanidine HCl.

Samples were analyzed with the following luciferase-based kits following the standard protocols: ATP Determination Kit (Invitrogen, A22066), ADP-Glo™ Kinase Assay (Promega,V6930), and GTPase-Glo™ Assay (Promega, V7681). Standards of nucleotides were made using nucleotides provided in the kits and diluting in the following buffer that matches the final buffer of the samples: 25 mM HEPES pH 7.5, 200 mM NaCl, 10 mM $MgCl_2$, 60 mM guanidine HCl. Standard concentrations were 1000 nM, 500 nM, 250 nM, 125 nM, 62.5 nM, 31.25 nM, 15.625 nM, and 7.8125 nM and were run with every replicate. Luminescence of samples was measured with a BioTek Synergy H1 hybrid reader at room temperature.Linear regressions of the standards were used to determine concentrations of samples after accounting for dilution.

This assay was validated by comparing the nucleotide measurements in the poly-L-Lysine dense and dilute phases with nucleotide measurements from absorbance values at 260 nm of the guanidine diluted samples. Poly-L-Lysine was prepared and measurements were performed as previously described(*40*). Using the Abs260, we calculated the concentration of nucleotide from the following extinction coefficients: ATP 15,400 $M^{-1}cm^{-1}$, ADP 15,400 $M^{-1}cm^{-1}$, GTP 11,700 $M^{-1}cm^{-1}$.

<u>BODIPY-ATP-γ-S localization in live cells</u>

Cells were transfected with mCherry-paxillin, as previously described. After 24 hours, cells were seeded onto fibronectin coated coverslips for 1.5 hours. Then cells were rinsed with buffer (10 mM MES pH 6.1, 138 mM KCl, 3 mM MgCl2, 2 mM EGTA). Cells were incubated in buffer containing 0.01% Saponin at room temperature for 1 min. Cells were gently rinsed in buffer and incubated with buffer containing 10 μM BODIPY or 10 μM BODIPY-ATP-γ-S for 5 min. Cells were quickly rinsed three times with buffer and imaged immediately. Images were acquired using TIRF microscopy within 5 minutes of rinsing. Pearson's correlation between mCherry-paxillin image and the BODIPY image was calculated in ImageJ.

Phase separation Prediction

The phase separation prediction scores (p-scores, 0-1) for human kinases were extracted from three published studies(*32-34*) and compiled into the supplemental spreadsheet (Data S1). For Fig. 5E, kinases with a p-score >0.7 from all three studies were classified as likely to phase separate (PS), while kinases with a p-score <0.3 from all three studies were classified as not likely to phase separate (No PS). GO term analysis for using pantherdb.org showed that PS kinases were slightly overrepresented (1.4 fold enrichment) with cytosol cellular component but had no significant difference in any other cellular component relative to the entire human kinome.

Protein expression and purification

*FAK and Abl*

His-mEGFP-FAK, His-mEGFP-FAK-W266A, FAK, FAK-W266A, and His-mEGFP-Abl1 were expressed from baculovirus in HighFive$^{TM}$ (Thermo) *Trichoplusia ni* cells. Cells were collected by centrifugation and lysed by douncing on ice in 25 mM HEPES (pH 7.5), 30 mM Imidazole (pH 7.5), 500 mM NaCl, 10% glycerol and 5 mM βME + cOmplete(TM), EDTA- free Protease Inhibitor tablet. Centrifugation- cleared lysate was applied to Ni- NTA agarose beads (Qiagen), washed with 25 mM HEPES (pH 7.5), 20 mM Imidazole (pH 7.5), 500 mM NaCl, 10% glycerol, 5 mM βME, 1 µg/ml benzamidine, and eluted with 25 mM HEPES (pH 7.5), 400 mM Imidazole (pH 7.5), 1 M NaCl, 10% glycerol, 5 mM βME, 1 µg/ml benzamidine. Protein was further purified with size exclusion chromatography using a Superdex 200 column (Cytiva) in 25 mM HEPES (pH 7.5), 500 mM NaCl, 10% Glycerol and 1 mM DTT. The His and GFP tags were optionally cleaved at this step using TEV protease treatment for 2 hr at room temperature. Proteins were optionally dephosphorylated at this step by addition of 2x molar equivalents of GST-PTP1B and incubating for 2 hr at room temperature. If TEV treatment and dephosphorylation were both done they were done at the same time. GST-PTP1B was removed at this step by supplementing protein with 20 mM imidazole and applying to Glutathione Sepharose 4B (Cytiva) resin and collecting the flowthrough. TEV treated or dephophorylated rotein was further purified with a second round of size exclusion chromatography.

*GST-PTP1B*

BL21(DE3) cells expressing GST- PTP1B were collected by centrifugation and lysed by sonication in 20 mM Tris- HCl (pH 8.0), 200 mM NaCl, 2 mM EDTA (pH 8.0), 1 mM DTT, 1 mM PMSF, 1 µg/ml antipain, 1 µg/ml benzamidine, 1 µg/ml leupeptin, and 1 µg/ml pepstatin. Centrifugation- cleared lysate was applied to Glutathione Sepharose 4B (Cytiva) and washed with 25 mM Tris- HCl (pH 8.0), 200 mM NaCl, and 1 mM DTT. Protein was eluted with 10 mM L-Glutathione (pH 8.0), 50 mM Tris HCL (pH 8.0), 200 mM NaCl, 1mM DTT. Protein was applied to a Source 15 Q anion exchange column and eluted with a gradient of 0 → 1000 mM NaCl in 20 mM HEPES (pH 7.5) and 1 mM DTT. Eluted protein was concentrated using an Amicon Ultra 10 k concentrator and further purified by size exclusion chromatography to remove any protein aggregates using a Superdex 75 prepgrade column (Cytiva) in 50 mM HEPES (pH 7.5), 150 mM NaCl, and 1 mM DTT.

*IDRs*

FAK IDR, Abl IDR, mEGFP-DDX21-CIDR-WT and mEGFP-DDX21-CIDR-R/S proteins were all expressed in Rosetta(DE3) cells with a C-terminal His-SUMO tag. Cells expressing the proteins were collected by centrifugation and lysed by cell disruption (Emulsiflex- C5, Avestin) in 20 mM Tris- HCl (pH 8.0), 300 mM NaCl, 40 mM Imidazole (pH 8.0), 10% glycerol, 1 mM BME, 1 μg/ml antipain, 1 μg/ml benzamidine, 1 μg/ml leupeptin, and 1 μg/ml pepstatin. Centrifugation-cleared lysate was applied to Ni- NTA agarose (Qiagen) and washed with 20 mM Tris- HCl (pH 8.0), 300 mM NaCl, 40 mM Imidazole (pH 8.0), 10% glycerol, 1 mM BME, 1 μg/ml benzamidine and eluted with 20 mM Tris- HCl (pH 8.0), 300 mM NaCl, 300 mM Imidazole (pH 8.0), 10% glycerol, 1 mM BME, 1 μg/ml benzamidine. Protein was applied to a SOURCE™ 15Q anion exchange column (Cytiva) and eluted with a gradient of 0 → 1,000 mM NaCl in 20 mM Tris- HCl (pH 8.0), 10% glycerol and 1 mM DTT. Eluted protein was treated with Ulp1 protease to cleave the His-SUMO tag overnight at 4°C. 20 mM imidazole (pH 8.0) was added to the cleaved protein which was then applied to Ni- NTA agarose (Qiagen) and the flowthrough was collected. This was concentrated using Amicon Ultra 10 k concentrators and further purified by size exclusion chromatography to remove any protein aggregates using a Superdex 200 column (Cytiva) in 25 mM HEPES (pH 7.5), 300 mM NaCl, 10% glycerol and 1 mM DTT. For the Abl IDR, the protein was applied to a SOURCE™ 15S cation exchange column instead of the SOURCE™ 15Q and eluted with a gradient of 0→ 1,000 mM NaCl in 25mM HEPES pH 7.5, 10% Glycerol, 1mM BME. The Abl IDR was also cleaved before applying to the SOURCE™ 15S cation exchange column.

*MST2*

MST2 protein was expressed in Rosetta(DE3) cells with a C-terminal His-SUMO tag. Cells expressing the protein were collected by centrifugation and lysed by cell disruption (Emulsiflex- C5, Avestin) in 20 mM Tris- HCl (pH 8.0), 300 mM NaCl, 40 mM Imidazole (pH 8.0), 10% glycerol, 1 mM BME, 1 μg/ml antipain, 1 μg/ml benzamidine, 1 μg/ml leupeptin, and 1 μg/ml pepstatin. Centrifugation-cleared lysate was applied to Ni- NTA agarose (Qiagen) and washed with 20 mM Tris- HCl (pH 8.0), 300 mM NaCl, 40 mM Imidazole (pH 8.0), 10% glycerol, 1 mM BME, 1 μg/ml benzamidine and eluted with 20 mM Tris- HCl (pH 8.0), 300 mM NaCl, 300 mM Imidazole (pH 8.0), 10% glycerol, 1 mM BME, 1 μg/ml benzamidine. Protein was applied to a Source 15 Q anion exchange column and eluted with a gradient of 0 → 1,000 mM NaCl in 20 mM Tris- HCl (pH 8.0), 10% glycerol and 1 mM DTT. Eluted protein was treated with Ulp1 protease to cleave the His-SUMO tag overnight at 4°C. 20 mM imidazole (pH 8.0) was added to the cleaved protein which was then applied to Ni- NTA agarose (Qiagen) and the flowthrough was collected. This was concentrated using Amicon Ultra 10 k concentrators and further purified by size exclusion chromatography to remove any protein aggregates using a Superdex 75 column (Cytiva) in 25 mM HEPES (pH 7.5), 300 mM NaCl, and 1 mM DTT.

*GFP-SAV1-MBP*

mEGFP-SAV1-MBP was expressed in Rosetta(DE3) cells. LB media was supplemented with 0.2% glucose. Cells expressing the protein were collected by centrifugation and lysed by cell disruption (Emulsiflex- C5, Avestin) in 20 mM Tris-HCl (pH 7.5), 200 mM NaCl, 1 mM EDTA, 1 mM DTT, 1 μg/ml antipain, 1 μg/ml benzamidine, 1 μg/ml leupeptin, and 1 μg/ml pepstatin.

Centrifugation-cleared lysate was applied to Amylose resin (NEB) and washed with 20 mM Tris-HCl (pH 7.5), 200 mM NaCl, 1 mM EDTA, 1 mM DTT, 1 µg/ml benzamidine and eluted with 20 mM Tris-HCl (pH 7.5), 200 mM NaCl, 10 mM maltose, 1 mM EDTA, 1 mM DTT, 1 µg/ml benzamidine. This was concentrated using Amicon Ultra 10 k concentrators and further purified by size exclusion chromatography to remove any protein aggregates using a Superdex 200 column (Cytiva) in 25 mM HEPES (pH 7.5), 300 mM NaCl, and 1 mM DTT.

*p130Cas, Paxillin, Nck, and N-WASP* were purified as previously described (*24*).

Fluorophore conjugation

For conjugation with maleimide chemistry, recombinant proteins to be labeled with Alexa fluorophores were concentrated using Amicon Ultra Centrifugal Filter units (Millipore) to ~100 µM. 5 mM βME was added to reduce cysteine residues followed by buffer exchange using a HiTrap 26/10 Desalting column (Cytiva) in protein storage buffer (see protein purification protocol) without any reducing agent. Fractions containing protein were collected and concentrated to 100 µM. 500 µM Alexa Fluor 647$C_2$ Maleimide (ThermoFisher) was added, and the reaction was incubated at 4 °C for 16 hr. The reaction was quenched with 1 µl 14.3 M βME followed by final buffer exchange using a HiTrap 26/10 Desalting column (Cytiva) in protein storage buffer buffer (see protein purification protocol) with reducing agent. Final protein concentration and degree of labeling were calculated from the protein absorbance using the following formula:

Concentration (M) = $A_{280}$ – ($A_{650}$ × 0.03)/ExtCoef

Degree of labeling = $A_{650}$/(239,000 x [protein conc.])

Visualization

The PhotoMol webtool was used to graph normalized counts of mass photometry data for each sample[3].

Diagrams of FAK, Abl and DDX21 domains and IDRs were made with IBS 2.0 webtool[4].

Dendrogram of human kinome was made with KinMap[5].
Diagrams of experimental setups (Fig. 1E, Fig. 2A, Fig. 4B) were created in BioRender. Case, L. (2025) https://BioRender.comhttps://biorender.com/phz7td7

### Supplementary Text

Explanation of volume-normalized fold rate enhancement of the dense phase

The percent of the total volume occupied by the dense phase (ie. droplet volume fraction; DVF) was calculated from the data in Fig. 1H and Fig. 1I using the equation $DVF = \frac{[Total] - [Dilute]}{[Dense] - [Dilute]}$ (see derivation below). Since the experiments in Fig. 1G used identical conditions and total FAK concentrations, this DVF was used to determine the volume-normalized fold rate enhancement of the dense phase ($VNFRE_{dense}$) using the equation $VNFRE_{dense} = \frac{(\frac{Dense\ rate}{DVF})}{(\frac{Dilute\ rate}{(1-DVF)})}$. The $VNFRE_{dense}$ was calculated for each of the 3 replicates in Fig. 1G generating N=3 values plotted in Fig. 1J. The dilute rate was measured directly, while the dense rate was determined by subtracting the dilute rate from the total rate.

Derivation of $DVF = \frac{[Total] - [Dilute]}{[Dense] - [Dilute]}$

$V_{Den}$ = volume of dense phase
$V_{Dil}$ = volume of dilute phase
$V_{Tot}$ = total solution volume
$[Dense]$ = dense phase concentration
$[Dilute]$ = dilute phase concentration
$[Total]$ = total solution concentration
$DVF$ = droplet volume fraction; percent of total solution occupied by dense phase

By definition:

$Eq. 1$ $$DVF = \frac{V_{Den}}{V_{Tot}}$$

$Eq. 2$ $$V_{Tot} = V_{Den} + V_{Dil}$$

Dividing each side of *Eq. 2* by $V_{Tot}$ yields:

$$1 = DVF + \frac{V_{Dil}}{V_{Tot}}$$

Subtracting $DVF$ from each side yields:

$Eq. 3$ $$1 - DVF = \frac{V_{Dil}}{V_{Tot}}$$

By conservation of mass:

$$(V_{Den})[Dense] + (V_{Dil})[Dilute] = (V_{Tot})[Total]$$

Dividing each side by $(V_{Tot})$ and substituting in *Eq. 1* and *Eq. 3* yields:

$$DVF[Dense] + (1 - DVF)[Dilute] = [Total]$$

Distributing $[Dilute]$ yields:

$$DVF[Dense] + [Dilute] - (DVF)[Dilute] = [Total]$$

Subtracting $[Dilute]$ from both sides and factoring out $DVF$ on the left side yields

$$DVF([Dense] - [Dilute]) = [Total] - [Dilute]$$

Finally, dividing both sides by ($[Dense] - [Dilute]$) yields

$$DVF = \frac{[Total] \; - \; [Dilute]}{[Dense] - [Dilute]}$$

Using our measured values of $[Dilute]$ = 0.0452 $\mu$M and $[Dense]$ = 106 $\mu$M, and knowing our total concentration of FAK added in each experiment (1 $\mu$M) the DVF is calculated to be 0.9%. We believe this to be a conservative upper bound of the DVF under these conditions since our measured $[Dense]$ value is likely smaller than the actual value. This is because under these conditions FAK forms relatively small (<5 $\mu$m) droplets which likely yield arbitrarily smaller mean intensity values due to the point spread function. Therefore, we also believe our calculated $VNFRE_{dense}$ values are conservative lower bound estimates, since a lower DVF values would yield a larger $VNFRE_{dense}$ calculation.

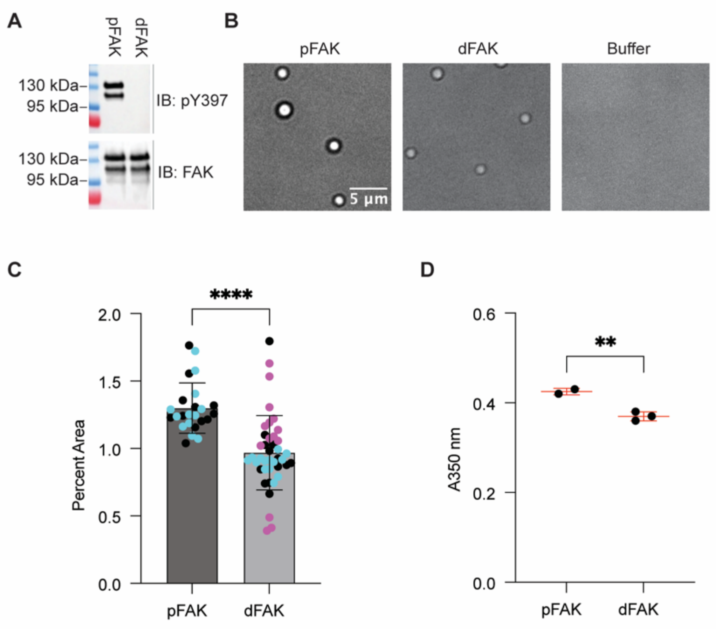


**Fig. S1. Tyrosine phosphorylation is not required for FAK phase separation.** (**A**) Chemiluminescent western blots against total and phospho-Y397 FAK of tyrosine dephosphorylated (dFAK) and native phosphorylated FAK (pFAK). (**B**) DIC microscopy images of pFAK and dFAK condensates at 1 µM and buffer only control. (**C**) Quantification of percent droplet area for data in (A). Each point represents a single field of view. N=24 for pFAK and N=41 for dFAK. Colors correspond to field of views from the same well. (**D**) Turbidity measurements of pFAK and dFAK at 2 µM. N=2 for pFAK and N=3 for dFAK. For all graphs error bars are standard deviation and significance was tested with unpaired T-tests. (* $p<0.0332$, ** $p<0.0021$, *** $p<0.0002$, **** $p<0.0001$). For all experiments the buffer was 25 mM HEPES pH 7.5, 50 mM NaCl, 1% glycerol, 1 mM DTT.

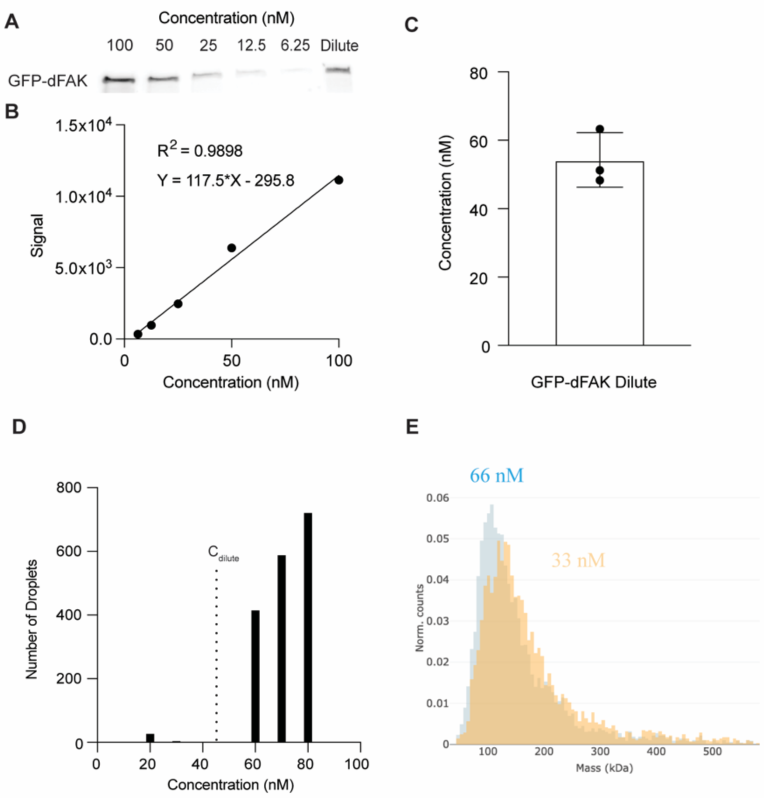


**Fig. S2. FAK condensates form above the measured $C_{dilute}$ in vitro with no detectable change in dilute phase oligomerization.** (**A**) Representative quantitative western blot of mEGFP-dFAK standards and dilute phase sample after sedimentation. (**B**) Quantification of bands in (A) and associated linear regression. (**C**) Quantification of mEGFP-dFAK dilute phase concentration measured from quantitative western blots (N=3 replicates). Error bars are standard deviation. (**D**) Quantification of number of droplets across 10 fields of view for 10, 20, 30, 60, 70, and 80 nM GFP-dFAK. (**E**) Mass photometry measurements of mEGFP-FAK protein in dilute phase below (orange; 33 nM) and above (blue; 66 nM) the $C_{dilute}$ concentration. For all experiments the buffer was 25 mM HEPES pH 7.5, 50 mM NaCl, 1% glycerol, 1 mM DTT.

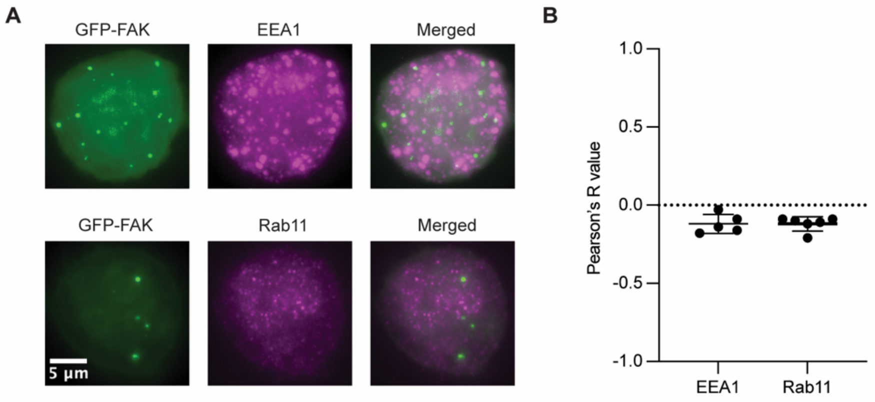


**Fig S3. mEGFP-FAK cytoplasmic puncta do not co-localize with early or recycling endosomes.** (**A**) Immunostaining of early (EEA1) and recycling (Rab11) endosomal markers in mEGFP-FAK-WT expressing cells plated on poly-D-Lysine. (**B**) Colocalization analysis of data in (A). Colocalization analysis was performed on mEGFP-FAK puncta and EEA1 and Rab11 endosomes for N=5 and N=6 immunostained cells, respectively. Analysis was limited to cells containing mEGFP-FAK puncta. Error bars represent standard deviation.

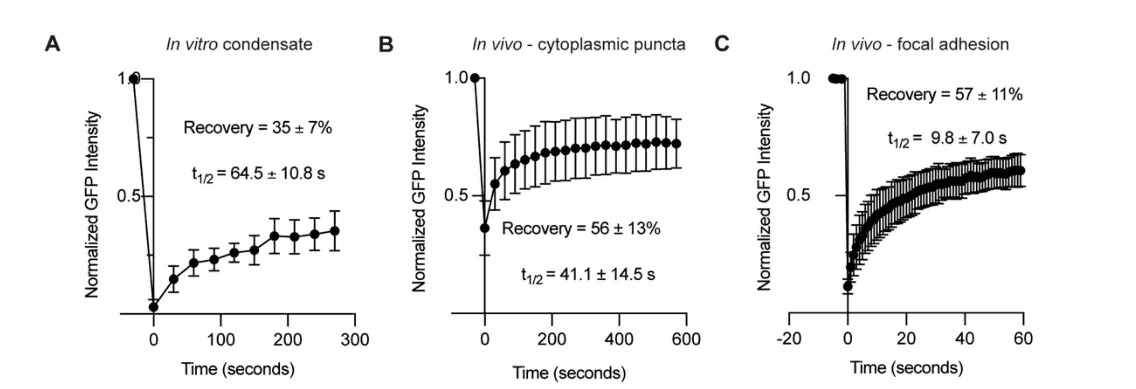


**Fig S4. mEGFP-FAK displays faster and higher FRAP recovery in cytoplasmic puncta and focal adhesions compared to in vitro condensates.** (**A**) FRAP data of mEGFP-FAK condensates in vitro. Buffer was 25 mM HEPES pH 7.5, 50 mM NaCl, 1% glycerol, 1 mM DTT (N = 4 replicates). (**B**) FRAP data of cytoplasmic mEGFP-FAK puncta (Identical data from Fig. 2C placed here for reference). All measurements were taken within 30 minutes of plating on poly-D-Lysine (N=5 replicates). (**C**) FRAP data of mEGFP-FAK at focal adhesions. All measurements were taken 2 hours after plating on fibronectin (N=6 replicates). For all plots, error bars represent standard deviation and error of recovery and $t_{1/2}$ are mean and standard deviations.

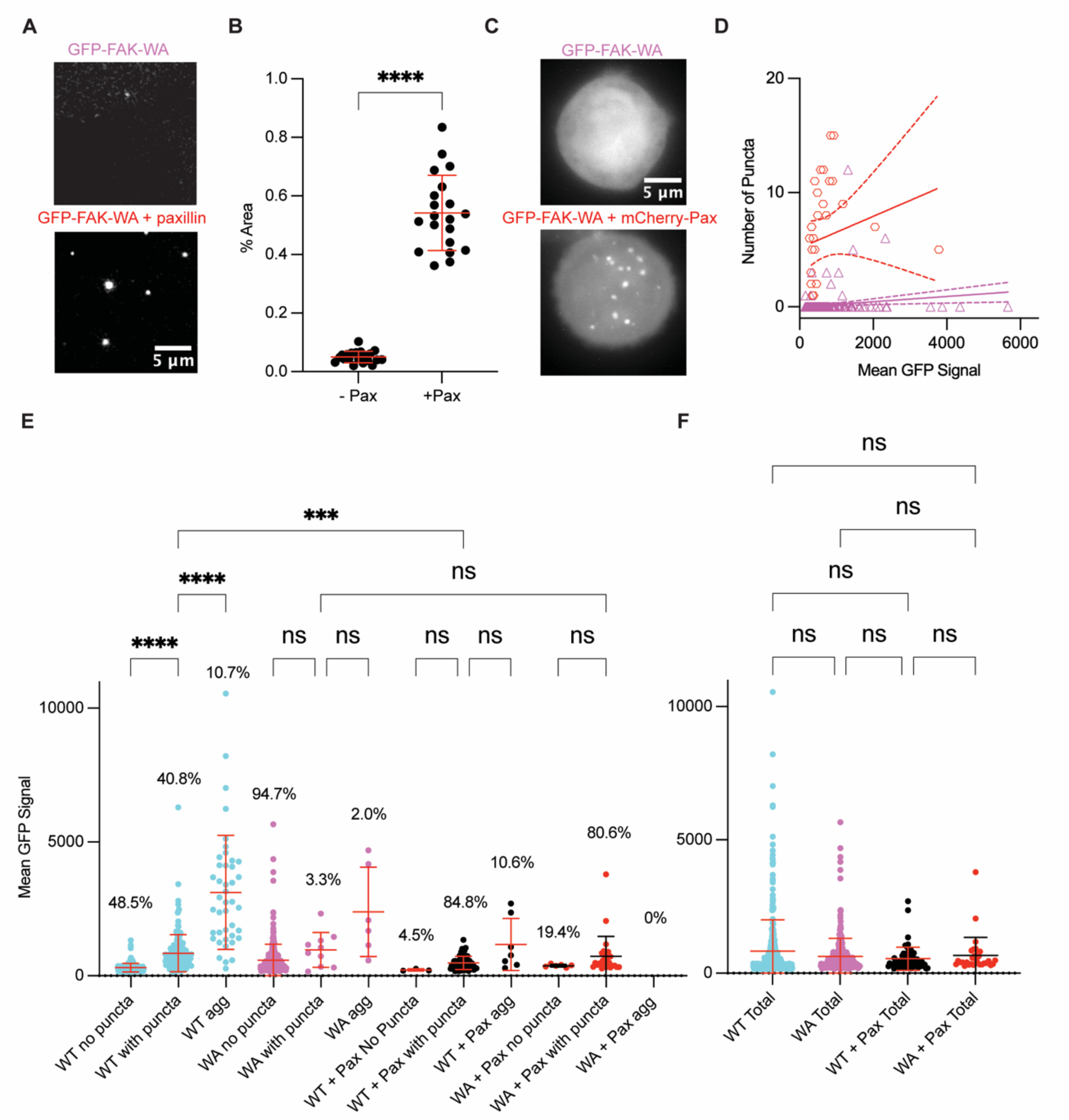


**Fig. S5. mEGFP-FAK puncta phenotypes in cells recapitulates in vitro treatments.** (**A**) Representative images of mEGFP-FAK-WA and mEGFP-FAK-WA + paxillin condensates in vitro. Final concentration of mEGFP-FAK and paxillin was 0.5 µM. Buffer: 25 mM HEPES pH 7.5, 50 mM NaCl, 1% glycerol, 1 mM DTT. (**B**) Quantification of percent area of condensates for both conditions in (A). Data is from 2 replicates (N=20 images). Significance was tested with unpaired t-test with Welch's correction. (**C**) Representative maximum intensity projections of MEFs plated on poly-D-Lysine. Cells chosen have a GFP signal of ~1000 a.u. (**D**) Quantification of number of puncta vs. total GFP signal. For cells co-expressed with paxillin, analyzed cells had an mCherry-Pax signal between 300-600 a.u. Dotted lines represent 95% confidence intervals of linear regressions. (**E**) Mean GFP signal of cells binned by puncta phenotypes (ie. no puncta, puncta, or aggregates). Percentages are the percent of transfected cells with each phenotype for each condition. (**F**) Mean GFP signal of all transfected cells for each condition. For (E) and (F)

significance was tested by Kruskal-Wallis test followed by a Dunn's multiple comparison test. For all graphs error bars represent standard deviation. * $p<0.0332$, ** $p<0.0021$, *** $p<0.0002$, **** $p<0.0001$

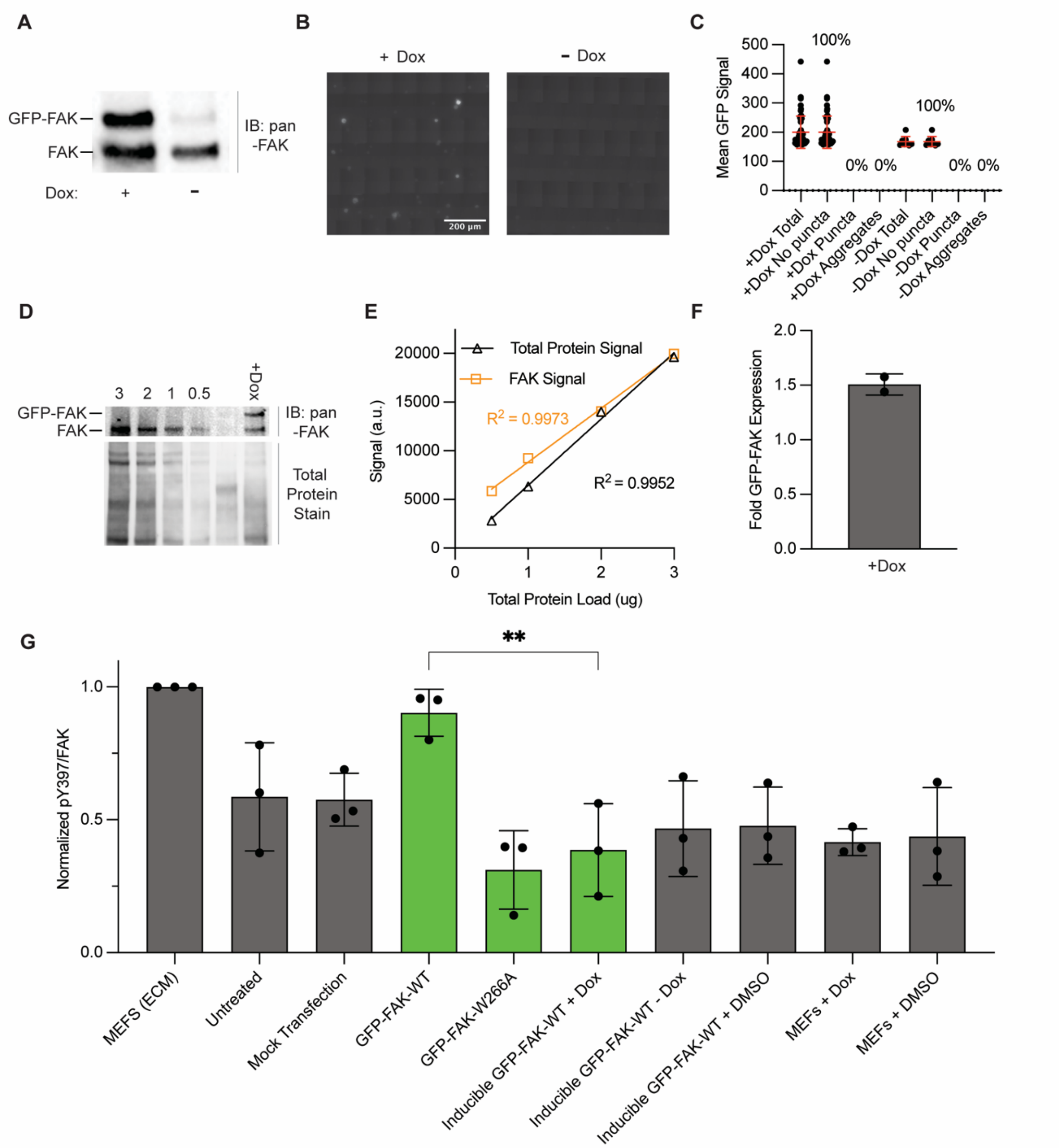


**Fig. S6. mEGFP-FAK overexpression is not sufficient to rescue integrin-dependent pY397/FAK ratios when plated on poly-D-Lysine.** (**A**) Representative western blot of stable Dox-inducible mEGFP-FAK MEF lysates with and without doxycycline treatment. (**B**) Images of stable Dox-inducible mEGFP-FAK MEF cells plated on poly-D-Lysine with and without doxycycline treatment. (**C**) Mean GFP signal for total populations of transfected MEFs or binned by puncta phenotypes (i.e.. no puncta, puncta, or aggregates). (**D**) Western blot and total protein stain of cell lysates from untransfected MEFs or Dox-inducible mEGFGP-FAK MEFs incubated with doxycycline for 24 hours. (**E**) Linearity of FAK channel and total protein stain was confirmed with dilutions of lysates. (**F**) Quantification of fold mEGFP-FAK expression from western blot in (D) for N=2 replicates. FAK was normalized to total protein stain and values represent fold expression compared to MEFs. (**G**) pY397/FAK ratios determined by western blot analysis of cell lysates. All cells were plated on poly-D-Lysine for 30 minutes before lysis except MEFS (ECM), which were grown on culture treated dishes for 24 hours. Green bars indicate analysis performed on mEGFP-FAK and grey bars on endogenous FAK. Significance tested by one-way ANOVA followed by a Tukey multiple comparison test (* $p<0.0332$, ** $p<0.0021$, *** $p<0.0002$, **** $p<0.0001$). For all graphs error bars represent standard deviation.

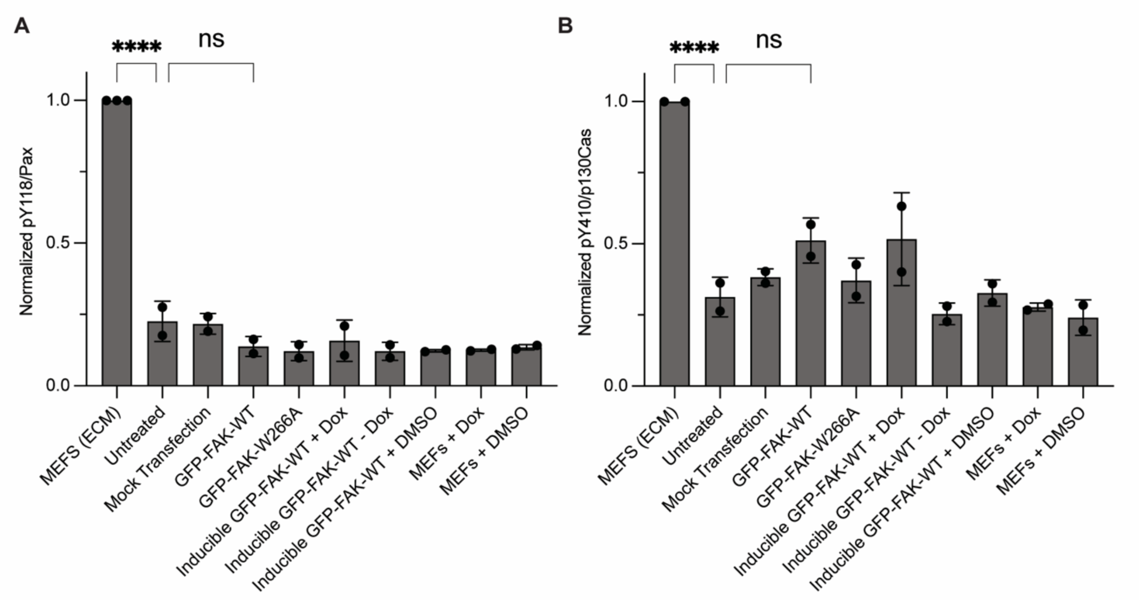


**Fig. S7. Cytoplasmic FAK condensates are not sufficient to rescue paxillin or p130Cas phosphorylation independent of integrin-based adhesion.** (**A**) pY118 Pax/Pax ratios determined by western blot analysis of MEF lysates. N=2 replicates. (**B**) pY410 Cas/p130Cas ratios determined by western blot analysis of MEF lysates. N=2 replicates. All cells were plated on poly-D-Lysine for 30 minutes before lysis except MEFS (ECM), which were grown on culture treated dishes for 24 hours. Significance tested by one-way ANOVA followed by a Tukey multiple comparison test (* $p<0.0332$, ** $p<0.0021$, *** $p<0.0002$, **** $p<0.0001$). For all graphs error bars represent standard deviation.

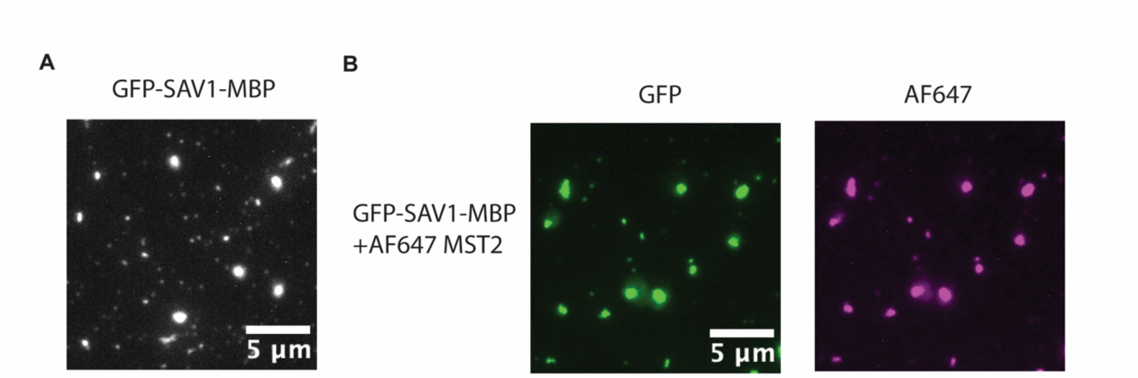


**Fig. S8. mEGFP-SAV1-MBP forms condensates that enrich Mst2 in vitro.** (**A**) Image of GFP-Sav1-MBP at 100 nM concentration. Final buffer composition is 25 mM HEPES pH 7.5, 100 mM NaCl, 10% PEG8000, 1 mM DTT. (**B**) Images of GFP-Sav1-MBP and AF647-Mst2 at 1 µM concentration. Final buffer composition is identical to (A).

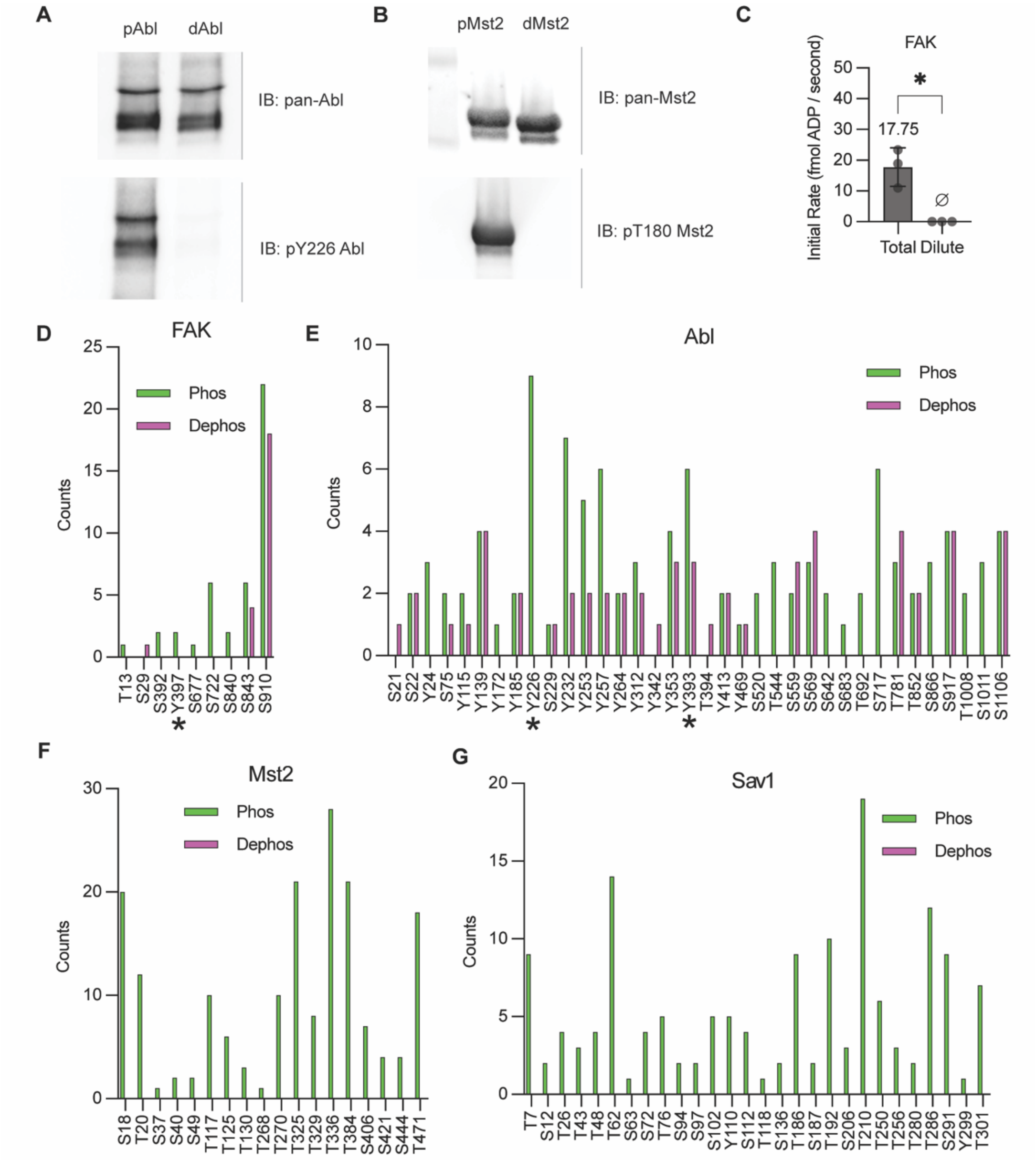


**Fig. S9. FAK, Abl and Mst2 undergo autophosphorylation on multiple sites in condensates.** (**A-B**) Representative western blots of Abl (A) and Mst2 (B) before and after autophosphorylation reactions. (**C**) Initial rates of FAK phosphorylation assays. Assays were performed in 25 mM HEPES pH 7.5, 50 mM NaCl, 1% glycerol (v/v), 1 mM DTT, 0.5 mM $MgCl_2$ and initiated by addition of 1 μM ATP. Error bars are standard deviation. N=3 replicates. Number above error bars is the mean. ø symbol denotes activity was undetectable. Significance was tested with unpaired t-test with Welch's correction (* p<0.0332, ** p<0.0021, *** p<0.0002, **** p<0.0001). (**D-G**) Phospho-mass spec results of before (dephos) and after (phos) phosphorylation reactions performed on total phase separated kinase solutions. Asterisks denotes canonical autophosphorylation sites where detected.

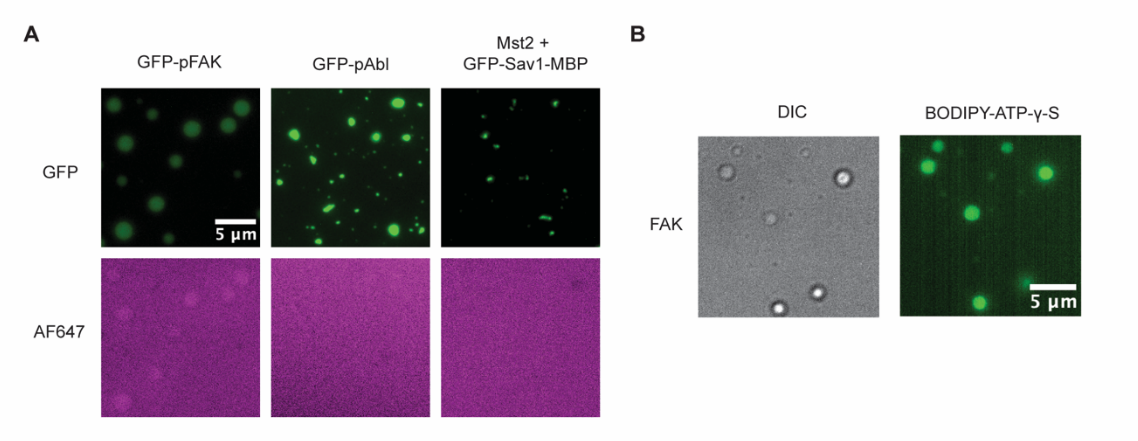


**Fig. S10. Kinase condensates enrich ATP independent of dye moiety.** (**A**) Microscopy images of 2.5 µM AF647 dye incubated with 1 µM mEGFP-FAK, 1 µM mEGFP-Abl or 100 nM mEGFP-Sav1-MBP + 100 nM Mst2. Final buffer conditions match those in Fig. 3A (Abl and Sav1+Mst2) and Fig. 1A (FAK). (**B**) Microscopy images of 1 µM FAK. Final buffer conditions match those in (A) except supplemented with 2.5 µM BODIPY-ATP-γ-S

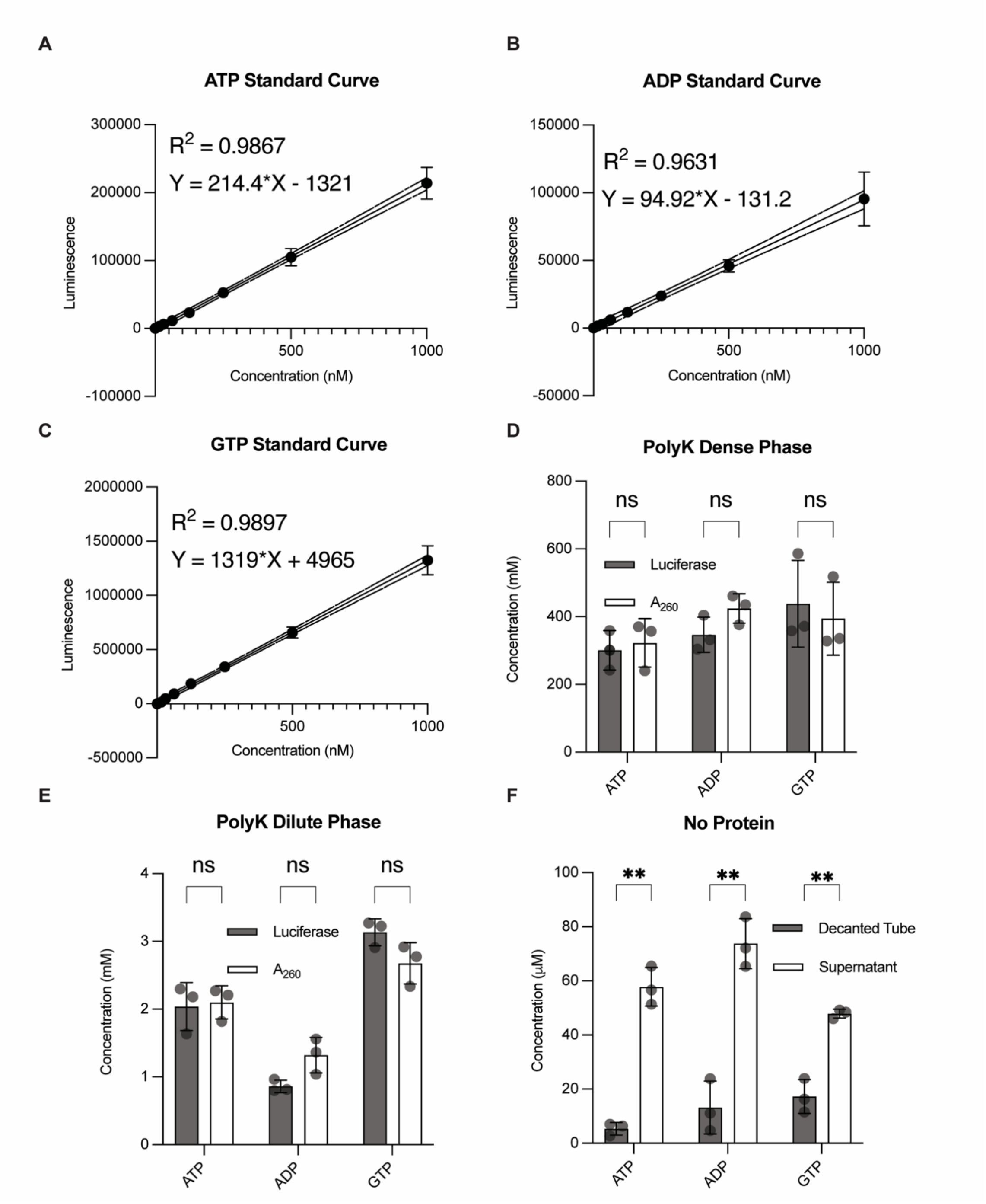


**Fig. S11. Condensate sedimentation-luciferase assays accurately measure condensate nucleotide concentrations.** (**A-C**) Standard curve of ATP (A), ADP (B), and GTP (C) standards in Nucleotide-Glo Assay Standard Buffer (25 mM HEPES pH 7.5, 200mM NaCl, 10 mM MgCl2, 60 mM guanidine HCl). For (A), (B), and (C) dotted lines represent 95% confidence intervals of linear regression. (**D-E**) Comparison of dense phase (D) and dilute phase (E) nucleotide concentrations measured from A260 spectrophotometry or sedimentation-luciferase assays of poly-L-Lysine condensates. Condensates were formed from ~100 µM poly-L-Lysine and 5 mM nucleotide in 10 mM immidazole pH 7.0 buffer. (**F**) Comparison of sedimentation-luciferase nucleotide measurements in solutions containing no protein. Samples taken from the supernatant and from resuspended decanted tube (capturing any nucleotides adsorbing to the tube). For all plots error bars represent standard deviation and statistical comparisons are unpaired t-tests with Welch correction (* $p<0.0332$, ** $p<0.0021$, *** $p<0.0002$, **** $p<0.0001$).

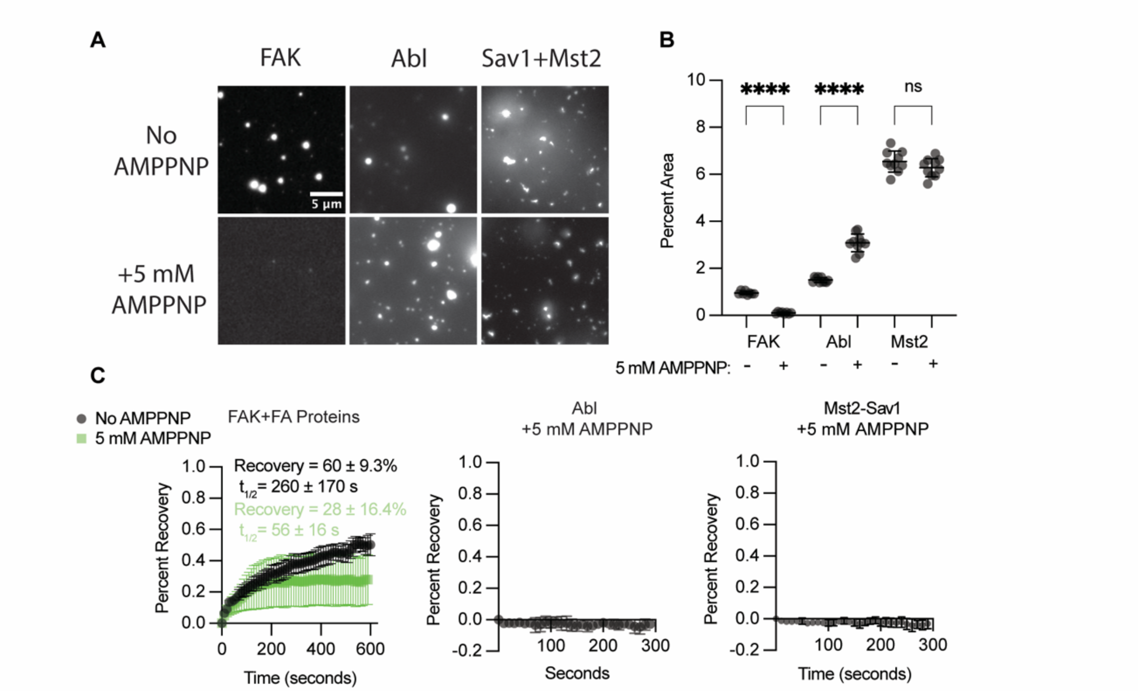


**Fig. S12. Physiological ATP concentrations do not inhibit Abl or Mst2 condensate formation and does not increase FRAP recovery of kinase condensates.** (**A**) Images of kinases with or without 5 mM AMPPNP. Buffers match those in Fig. 3B (Abl and Sav1+Mst2) and Fig.1A (FAK). (**B**) Percent area occupied by condensates for conditions in (A). Significance was tested with unpaired t-tests with Welch's correction (* p <0.0332, ** p<0.0021, *** p<0.0002, **** p<0.0001)(N>8 images for each condition)(**C**) FRAP analysis of kinase condensates with (black) or without (green) 5 mM AMPPNP. For FAK, the FA proteins Paxillin, Nck, N-WASP, and p130Cas where added at 1 µM. All conditions used buffers identical to those used in (A). For all graphs, error bars represent standard deviation. Error bars not shown are smaller than symbol.

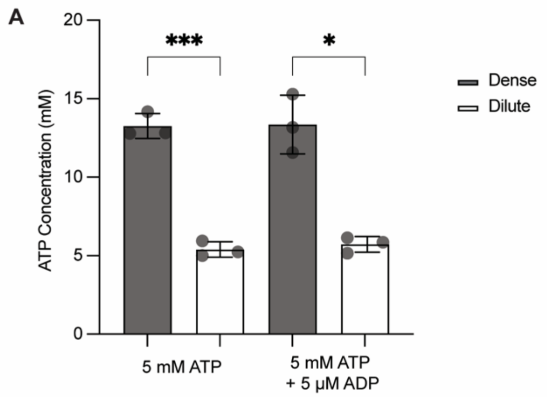


**Fig. S13. Reconstituted focal adhesion condensates enrich ATP at physiological ATP and ADP concentrations.** (**A**) Dense and dilute phase ATP concentration measurements of reconstituted focal adhesion condensates. FAK, paxillin, Nck, N-WASP and p130Cas are all 1 µM final concentration. Buffer: 25 mM HEPES pH 7.5, 50 mM NaCl, 1% glycerol (v/v), 1 mM DTT, 5 mM ATP with or without 5 µM ADP. N=3 replicates. Error bars denote standard deviation and statistical tests are unpaired T-tests with Welch correction (* $p<0.0332$, ** $p<0.0021$, *** $p<0.0002$, **** $p<0.0001$).

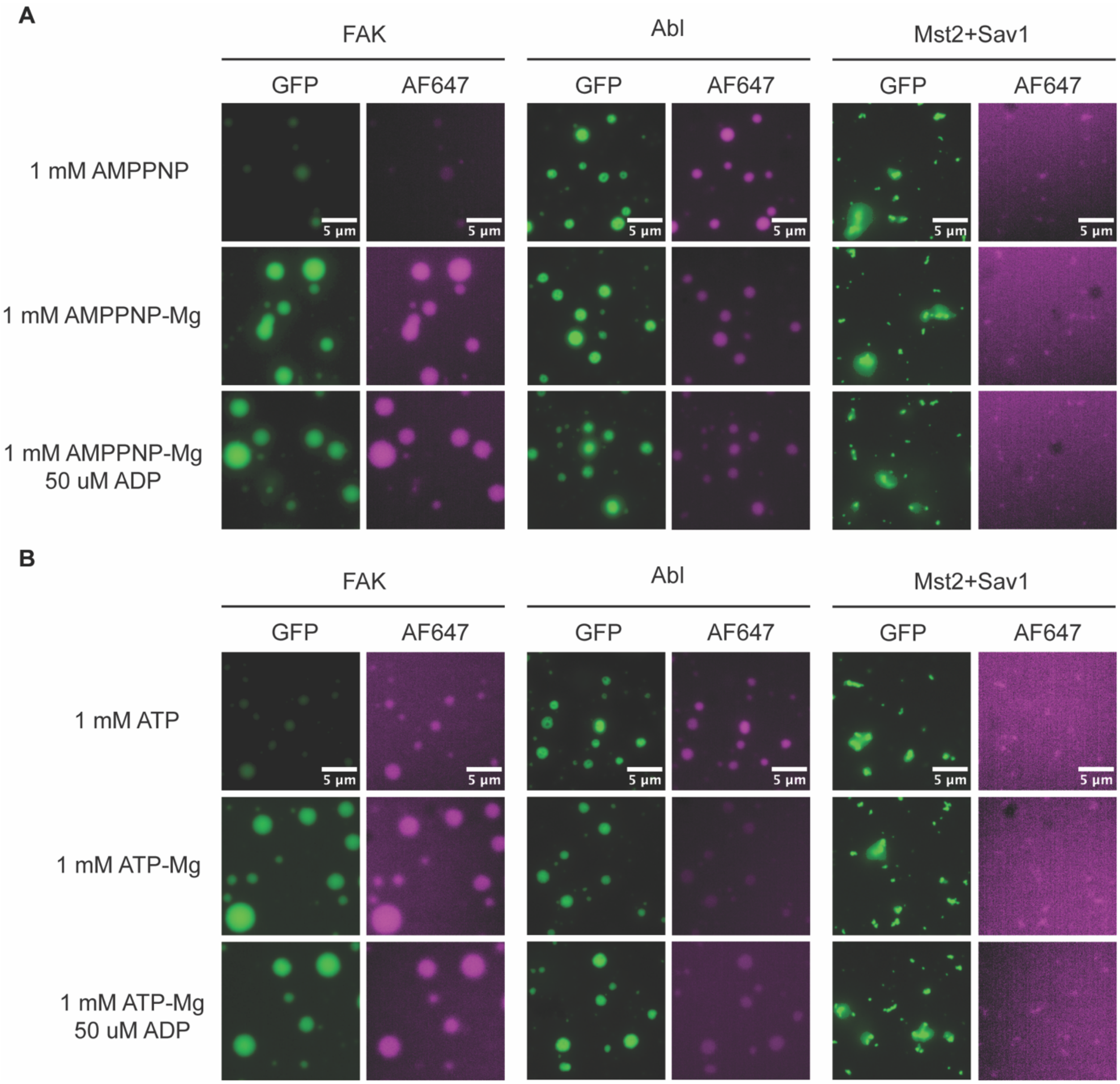


**Fig. S14. ATP enrichment in kinase condensates is not disrupted by the presence of physiological $Mg^{2+}$ or ADP concentrations.** (A) Representative fluorescent microscopy images of focal adhesion, mEGFP-Abl and Mst2+mEGFP-Sav1 condensates with 1mM AMPPNP with or without 1 mM $MgCl_2$ and 50 µM ADP. Final protein concentration is 1 µM for all conditions. Focal adhesion condensates were made from mEGFP-FAK, paxillin, Nck, N-WASP and p130Cas. Buffer: 25 mM HEPES pH 7.5, 100 mM NaCl, 10 µM AF647 ATP, 1% glycerol (v/v), 1 mM DTT supplemented with 5 or 10% PEG8000 for Abl and Mst2+Sav1, respectively. (**B**) Identical conditions in (A) except with ATP instead of AMPPNP.

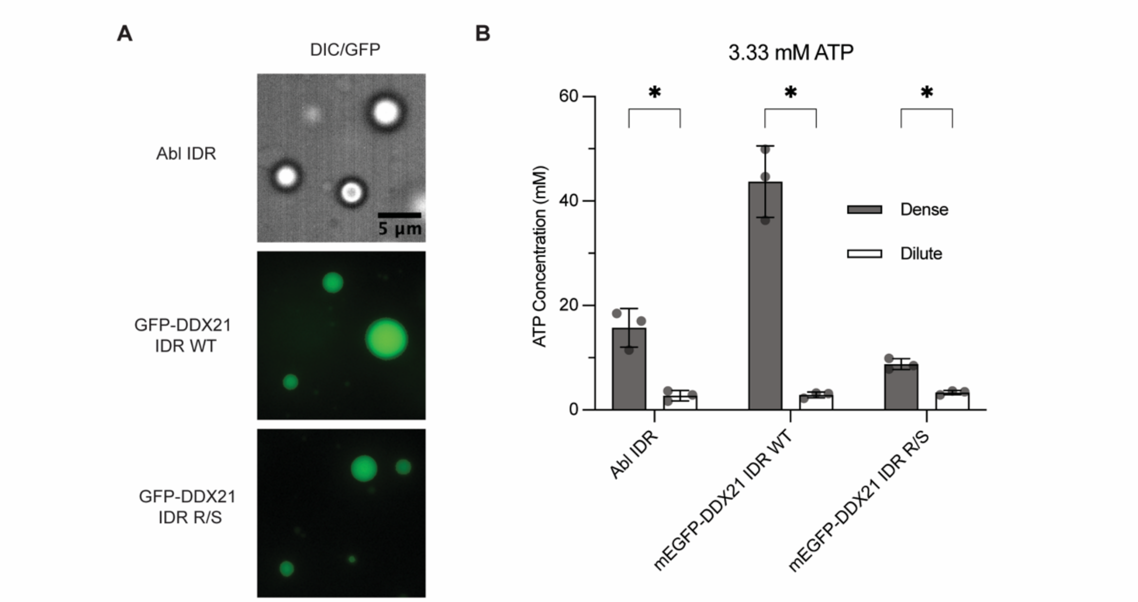


**Fig. S15. Abl and DDX21 IDR condensates enrich ATP at physiological ATP concentrations.** (**A**) Fluorescence (green) or DIC (greyscale) images of IDR condensates. Buffer conditions are 20 µM protein, 25 mM HEPES pH 7.5, 100 mM NaCl, 20% PEG8000, 1 mM DTT and 3.33 mM ATP. (**B**) Dense and dilute phase ATP concentration measurements from sedimentation-luciferase assays. Buffer conditions are identical to those in (A). N=3 replicates. Error bars denote standard deviation. Statistical tests are unpaired T-tests with Welch correction (* $p<0.0332$, ** $p<0.0021$, *** $p<0.0002$, **** $p<0.0001$).

| Protein | Species | Sequence |
|---|---|---|
| His-TEV-mEGFP-TEV-FAK-WT | Mouse (*Mus musculus*) | MSYYHHHHHHDYDIPTTENLYFQGAMGASMVSKGEELFTGVVPILVELD<br>GDVNGHKFSVSGEGEGDATYGKLTLKFICTTGKLPVPWPTLVTTLTYGVQC<br>FSRYPDHMKQHDFFKSAMPEGYVQERTIFFKDDGNYKTRAEVKFEGDTLV<br>NRIELKGIDFKEDGNILGHKLEYNYNSHNVYIMADKQKNGIKVNFKIRHNIE<br>DGSVQLADHYQQNTPIGDGPVLLPDNHYLSTQSKLSKDPNEKRDHMVLL<br>EFVTAAGITLGMDELYKGGSGGSENLYFQGGIRMAAAYLDPNLNHTPSSS<br>TKTHLGTGMERSPGAMERVLKVFHHFESSSEPTTWASIIRHGDATDVRGII<br>QKIVDSHKVKHVACYGFRLSHLRSEEVHWLHVDMGVSSVREKYELAHPPE<br>EWKYELRIRYLPKGFLNQFTEDKPTLNFFYQQVKSDYMQEIADQVDQEIAL<br>KLGCLEIRRSYWEMRGNALEKKSNYEVLEKDVGLKRFFPKSLLDSVKAKTLR<br>KLIQQTFRQFANLNREESILKFFEILSPVYRFDKECFKCALGSSWIISVELWIG<br>PEEGISYLTDKGCNPTHLADFNQVQTIQYSNSEDKDRKGMLQLKIAGAPEP<br>LTVTAPSLTIAENMADLIDGYCRLVNGATQSFIIRPQKEGERALPSIPKLANS<br>EKQGMRTHAVSVSETDDYAEIIDEEDTYTMPSTRDYEIQRERIELGRCIGEG<br>QFGDVHQGVYLSPENPALAVAIKTCKNCTSDSVREKFLQEALTMRQFDHP<br>HIVKLIGVITENPVWIIMELCTLGELRSFLQVRKYSLDLASLILYAYQLSTALA<br>YLESKRFVHRDIAARNVLVSSNDCVKLGDFGLSRYMEDSTYYKASKGKLPIK<br>WMAPESINFRRFTSASDVWMFGVCMWEILMHGVKPFQGVKNNDVIGR<br>IENGERLPMPPNCPPTLYSLMTKCWAYDPSRRPRFTELKAQLSTILEEEKV<br>QQEERMRMESRRQATVSWDSGGSDEAPPKPSRPGYPSPRSSEGFYPSPQ<br>HMVQTNHYQVSGYPGSHGIPAMAGSIYQGQASLLDQTELWNHRPQEM<br>SMWQPSVEDSAALDLRGMGQVLPPHLMEERLIRQQQEMEEDQRWLEK<br>EERFLKPDVRLSRGSIDREDGSFQGPTGNQHIYQPVGKPDPAAPPKKPPRP<br>GAPGHLSNLSSISSPADSYNEGVKLQPQEISPPPTANLDRSNDKVYENVTG<br>LVKAVIEMSSKIQPAPPEEYVPMVKEVGLALRTLLATVDETIPALPASTHREI<br>EMAQKLLNSDLGELISKMKLAQQYVMTSLQQEYKKQMLTAAHALAVDA<br>KNLLDVIDQARLKMLGQTRPH |
| His-TEV-mEGFP-TEV-FAK-W266A | Mouse (*Mus musculus*) | MSYYHHHHHHDYDIPTTENLYFQGAMGASMVSKGEELFTGVVPILVELD<br>GDVNGHKFSVSGEGEGDATYGKLTLKFICTTGKLPVPWPTLVTTLTYGVQC<br>FSRYPDHMKQHDFFKSAMPEGYVQERTIFFKDDGNYKTRAEVKFEGDTLV<br>NRIELKGIDFKEDGNILGHKLEYNYNSHNVYIMADKQKNGIKVNFKIRHNIE<br>DGSVQLADHYQQNTPIGDGPVLLPDNHYLSTQSKLSKDPNEKRDHMVLL<br>EFVTAAGITLGMDELYKGGSGGSENLYFQGGIRMAAAYLDPNLNHTPSSS<br>TKTHLGTGMERSPGAMERVLKVFHHFESSSEPTTWASIIRHGDATDVRGII<br>QKIVDSHKVKHVACYGFRLSHLRSEEVHWLHVDMGVSSVREKYELAHPPE<br>EWKYELRIRYLPKGFLNQFTEDKPTLNFFYQQVKSDYMQEIADQVDQEIAL<br>KLGCLEIRRSYWEMRGNALEKKSNYEVLEKDVGLKRFFPKSLLDSVKAKTLR<br>KLIQQTFRQFANLNREESILKFFEILSPVYRFDKECFKCALGSSAIISVELAIGP<br>EEGISYLTDKGCNPTHLADFNQVQTIQYSNSEDKDRKGMLQLKIAGAPEPL<br>TVTAPSLTIAENMADLIDGYCRLVNGATQSFIIRPQKEGERALPSIPKLANSE<br>KQGMRTHAVSVSETDDYAEIIDEEDTYTMPSTRDYEIQRERIELGRCIGEG |

| | | |
|---|---|---|
| | | QFGDVHQGVYLSPENPALAVAIKTCKNCTSDSVREKFLQEALTMRQFDHP HIVKLIGVITENPVWIIMELCTLGELRSFLQVRKYSLDLASLILYAYQLSTALA YLESKRFVHRDIAARNVLVSSNDCVKLGDFGLSRYMEDSTYYKASKGKLPIK WMAPESINFRRFTSASDVWMFGVCMWEILMHGVKPFQGVKNNDVIGR IENGERLPMPPNCPPTLYSLMTKCWAYDPSRRPRFTELKAQLSTILEEEKV QQEERMRMESRRQATVSWDSGGSDEAPPKPSRPGYPSPRSSEGFYPSPQ HMVQTNHYQVSGYPGSHGIPAMAGSIYQGQASLLDQTELWNHRPQEM SMWQPSVEDSAALDLRGMGQVLPPHLMEERLIRQQQEMEEDQRWLEK EERFLKPDVRLSRGSIDREDGSFQGPTGNQHIYQPVGKPDPAAPPKKPPRP GAPGHLSNLSSISSPADSYNEGVKLQPQEISPPPTANLDRSNDKVYENVTG LVKAVIEMSSKIQPAPPEEYVPMVKEVGLALRTLLATVDETIPALPASTHREI EMAQKLLNSDLGELISKMKLAQQYVMTSLQQEYKKQMLTAAHALAVDA KNLLDVIDQARLKMLGQTRPH |
| His-mEGFP-TEV-Abl1 | Human (*Homo sapiens*) | MSYYHHHHHHDYDIPTTAMGASMVSKGEELFTGVVPILVELDGDVNGHK FSVSGEGEGDATYGKLTLKFICTTGKLPVPWPTLVTTLTYGVQCFSRYPDH MKQHDFFKSAMPEGYVQERTIFFKDDGNYKTRAEVKFEGDTLVNRIELKG IDFKEDGNILGHKLEYNYNSHNVYIMADKQKNGIKVNFKIRHNIEDGSVQL ADHYQQNTPIGDGPVLLPDNHYLSTQSKLSKDPNEKRDHMVLLEFVTAAG ITLGMDELYKGGSGGSENLYFQGGIRMLEICLKLVGCKSKKGLSSSSSCYLEE ALQRPVASDFEPQGLSEAARWNSKENLLAGPSENDPNLFVALYDFVASGD NTLSITKGEKLRVLGYNHNGEWCEAQTKNGQGWVPSNYITPVNSLEKHS WYHGPVSRNAAEYLLSSGINGSFLVRESESSPGQRSISLRYEGRVYHYRINT ASDGKLYVSSESRFNTLAELVHHHSTVADGLITTLHYPAPKRNKPTVYGVSP NYDKWEMERTDITMKHKLGGGQYGEVYEGVWKKYSLTVAVKTLKEDTM EVEEFLKEAAVMKEIKHPNLVQLLGVCTREPPFYIITEFMTYGNLLDYLREC NRQEVNAVVLLYMATQISSAMEYLEKKNFIHRDLAARNCLVGENHLVKVA DFGLSRLMTGDTYTAHAGAKFPIKWTAPESLAYNKFSIKSDVWAFGVLLW EIATYGMSPYPGIDLSQVYELLEKDYRMERPEGCPEKVYELMRACWQWN PSDRPSFAEIHQAFETMFQESSISDEVEKELGKQGVRGAVSTLLQAPELPTK TRTSRRAAEHRDTTDVPEMPHSKGQGESDPLDHEPAVSPLLPRKERGPPE GGLNEDERLLPKDKKTNLFSALIKKKKKTAPTPPKRSSSFREMDGQPERRG AGEEEGRDISNGALAFTPLDTADPAKSPKPSNGAGVPNGALRESGGSGFR SPHLWKKSSTLTSSRLATGEEEGGGSSSKRFLRSCSASCVPHGAKDTEWRS VTLPRDLQSTGRQFDSSTFGGHKSEKPALPRKRAGENRSDQVTRGTVTPP PRLVKKNEEAADEVFKDIMESSPGSSPPNLTPKPLRRQVTVAPASGLPHKE EAGKGSALGTPAAAEPVTPTSKAGSGAPGGTSKGPAEESRVRRHKHSSES PGRDKGKLSRLKPAPPPPPAASAGKAGGKPSQSPSQEAAGEAVLGAKTKA TSLVDAVNSDAAKPSQPGEGLKKPVLPATPKPQSAKPSGTPISPAPVPSTLP SASSALAGDQPSSTAFIPLISTRVSLRKTRQPPERIASGAITKGVVLDSTEALC LAISRNSEQMASHSAVLEAGKNLYTFCVSYVDSIQQMRNKFAFREAINKLE NNLRELQICPATAGSGPAATQDFSKLLSSVKEISDIVQR |

| | | |
|---|---|---|
| His-<br>SUMO-<br>Mst2 | Human<br>(*Homo*<br>*sapiens*) | MGSSHHHHHHSSGLVPRGSHMASMSDSEVNQEAKPEVKPEVKPETHIN<br>LKVSDGSSEIFFKIKKTTPLRRLMEAFAKRQGKEMDSLRFLYDGIRIQADQT<br>PEDLDMEDNDIIEAHREQIGGSMEQPPAPKSKLKKLSEDSLTKQPEEVFDV<br>LEKLGEGSYGSVFKAIHKESGQVVAIKQVPVESDLQEIIKEISIMQQCDSPYV<br>VKYYGSYFKNTDLWIVMEYCGAGSVSDIIRLRNKTLIEDEIATILKSTLKGLEY<br>LHFMRKIHRDIKAGNILLNTEGHAKLADFGVAGQLTDTMAKRNTVIGTPF<br>WMAPEVIQEIGYNCVADIWSLGITSIEMAEGKPPYADIHPMRAIFMIPTN<br>PPPTFRKPELWSDDFTDFVKKCLVKNPEQRATATQLLQHPFIKNAKPVSILR<br>DLITEAMEIKAKRHEEQQRELEEEEENSDEDELDSHTMVKTSVESVGTMR<br>ATSTMSEGAQTMIEHNSTMLESDLGTMVINSEDEEEEDGTMKRNATSPQ<br>VQRPSFMDYFDKQDFKNKSHENCNQNMHEPFPMSKNVFPDNWKVPQ<br>DGDFDFLKNLSLEELQMRLKALDPMMEREIEELRQRYTAKRQPILDAMDA<br>K |
| mEGFP-<br>Sav1-<br>TEV-MBP | Human<br>(*Homo*<br>*sapiens*) | MGSSMVSKGEELFTGVVPILVELDGDVNGHKFSVSGEGEGDATYGKLTLK<br>FICTTGKLPVPWPTLVTTLTYGVQCFSRYPDHMKQHDFFKSAMPEGYVQE<br>RTIFFKDDGNYKTRAEVKFEGDTLVNRIELKGIDFKEDGNILGHKLEYNYNS<br>HNVYIMADKQKNGIKVNFKIRHNIEDGSVQLADHYQQNTPIGDGPVLLPD<br>NHYLSTQSKLSKDPNEKRDHMVLLEFVTAAGITLGMDELYKGSMLSRKKT<br>KNEVSKPAEVQGKYVKKETSPLLRNLMPSFIRHGPTIPRRTDICLPDSSPNA<br>FSTSGDVVSRNQSFLRTPIQRTPHEIMRRESNRLSAPSYLARSLADVPREYG<br>SSQSFVTEVSFAVENGDSGSRYYYSDNFFDGQRKRPLGDRAHEDYRYYEY<br>NHDLFQRMPQNQGRHASGIGRVAATSLGNLTNHGSEDLPLPPGWSVD<br>WTMRGRKYYIDHNTNTTHWSHPLEREGLPPGWERVESSEFGTYYVDHT<br>NKKAQYRHPCAPSVPRYDQPPPVTYQPQQTERNQSLLVPANPYHTAEIPD<br>WLQVYARAPVKYDHILKWELFQLADLDTYQGMLKLLFMKELEQIVKMYE<br>AYRQALLTELENRKQRQQWYAQQHGKNFLEENLYFQGMKIEEGKLVIWI<br>NGDKGYNGLAEVGKKFEKDTGIKVTVEHPDKLEEKFPQVAATGDGPDIIF<br>WAHDRFGGYAQSGLLAEITPDKAFQDKLYPFTWDAVRYNGKLIAYPIAVE<br>ALSLIYNKDLLPNPPKTWEEIPALDKELKAKGKSALMFNLQEPYFTWPLIAA<br>DGGYAFKYENGKYDIKDVGVDNAGAKAGLTFLVDLIKNKHMNADTDYSIA<br>EAAFNKGETAMTINGPWAWSNIDTSKVNYGVTVLPTFKGQPSKPFVGVL<br>SAGINAASPNKELAKEFLENYLLTDEGLEAVNKDKPLGAVALKSYEEELAKD<br>PRIAATMENAQKGEIMPNIPQMSAFWYAVRTAVINAASGRQTVDEALKD<br>AQT |
| GST-<br>paxillin | Human<br>(*Homo*<br>*sapiens*) | MSPILGYWKIKGLVQPTRLLLEYLEEKYEEHLYERDEGDKWRNKKFELGLEF<br>PNLPYYIDGDVKLTQSMAIIRYIADKHNMLGGCPKERAEISMLEGAVLDIRY<br>GVSRIAYSKDFETLKVDFLSKLPEMLKMFEDRLCHKTYLNGDHVTHPDFML<br>YDALDVVLYMDPMCLDAFPKLVCFKKRIEAIPQIDKYLKSSKYIAWPLQGW<br>QATFGGGDHPPKSDLVPRGSENLYFQGHMDDLDALLADLESTTSHISKRP<br>VFLSEETPYSYPTGNHTYQEIAVPPPVPPPPSSEALNGTILDPLDQWQPSGS<br>RFIHQQPQSSSPVYGSSAKTSSVSNPQDSVGSPCSRVGEEEHVYSFPNKQK<br>SAEPSPTVMSTSLGSNLSELDRLLLELNAVQHNPPGFPADEANSSPPLPGA<br>LSPLYGVPETNSPLGGKAGPLTKEKPKRNGGRGLEDVRPSVESLLDELESSV |

| | | |
|---|---|---|
| | | PSPVPAITVNQGEMSSPQRVTSTQQQTRISASSATRELDELMASLSDFKF<br>MAQGKTGSSSPPGGPPKPGSQLDSMLGSLQSDLNKLGVATVAKGVCGA<br>CKKPIAGQVVTAMGKTWHPEHFVCTHCQEEIGSRNFFERDGQPYCEKDY<br>HNLFSPRCYYCNGPILDKVVTALDRTWHPEHFFCAQCGAFFGPEGFHEKD<br>GKAYCRKDYFDMFAPKCGGCARAILENYISALNTLWHPECFVCRECFTPFV<br>NGSFFEHDGQPYCEVHYHERRGSLCSGCQKPITGRCITAMAKKFHPEHFV<br>CAFCLKQLNKGTFKEQNDKPYCQNCFLKLFC |
| GST-PTP1B | Human (*Homo sapiens*) | MSPILGYWKIKGLVQPTRLLLEYLEEKYEEHLYERDEGDKWRNKKFELGLEF<br>PNLPYYIDGDVKLTQSMAIIRYIADKHNMLGGCPKERAEISMLEGAVLDIRY<br>GVSRIAYSKDFETLKVDFLSKLPEMLKMFEDRLCHKTYLNGDHVTHPDFML<br>YDALDVVLYMDPMCLDAFPKLVCFKKRIEAIPQIDKYLKSSKYIAWPLQGW<br>QATFGGGDHPPKSDLVPRGSPEFPGRLERPHRDGSGVLFQGPMEMEKEF<br>EQIDKSGSWAAIYQDIRHEASDFPCRVAKLPKNKNRNRYRDVSPFDHSRIK<br>LHQEDNDYINASLIKMEEAQRSYILTQGPLPNTCGHFWEMVWEQKSRGV<br>VMLNRVMEKGSLKCAQYWPQKEEKEMIFEDTNLKLTLISEDIKSYYTVRQL<br>ELENLTTQETREILHFHYTTWPDFGVPESPASFLNFLFKVRESGSLSPEHGP<br>VVVHCSAGIGRSGTFCLADTCLLLMDKRKDPSSVDIKKVLLEMRKFRMGLI<br>QTADQLRFSYLAVIEGAKFIMGDSSVQDQWKELSHEDLEPPPEHIPPPPRP<br>PKRILEPHN |
| His-p130Cas | Rat (*Rattus norvegicus*) | MSYYHHHHHHDYDIPTTENLYFQGAMGSGTMKYLNVLAKALYDNVAES<br>PDELSFRKGDIMTVLERDTQGLDGWWLCSLHGRQGIVPGNRLKILVGMY<br>DKKPAAPGPGPPATPPQPQPSLPQGVHTPVPPASQYSPMLPTAYQPQPD<br>NVYLVPTPSKTQQGLYQAPGPNPQFQSPPAKQTSTFSKQTPHHSFPSPAT<br>DLYQVPPGPGSPAQDIYQVPPSAGTGHDIYQVPPSLDTRSWEGTKPPAKV<br>VVPTRVGQGYVYEASQAEQDEYDTPRHLLAPGSQDIYDVPPVRGLLPNQY<br>GQEVYDTPPMAVKGPNGRDPLLDVYDVPPSVEKGLPPSNHHSVYDVPPS<br>VSKDVPDGPLLREETYDVPPAFAKPKPFDPTRHPLILAAPPPDSPPAEDVYD<br>VPPPAPDLYDVPPGLRRPGPGTLYDVPRERVLPPEVADGSVIDDGVYAVP<br>PPAEREAPTDGKRLSASSTGSTRSSQSASSLEVVVPGREPLELEVAVETLAR<br>LQQGVSTTVAHLLDLVGSASGPGGWRSTSEPQEPPVQDLKAAVAAVHGA<br>VHELLEFARSAVSSATHTSDRTLHAKLSRQLQKMEDVYQTLVVHGQVLDS<br>GRGGPGFTLDDLDRLVACSRAVPEDAKQLASFLHGNASLLFRRTKAPGPG<br>PEGSSSLHLNPTDKASSIQSRPLPSPPKFTSQDSPDGQYENSEGGWMEDY<br>DYVHLQGKEEFEKTQKELLEKGNIVRQGKGQLELQQLKQFERLEQEVSRPI<br>DHDLANWTPAQPLVPGRTGGLGPSDRQLLLFYLEQCEANLTTLTDAVDAF<br>FTAVATNQPPKIFVAHSKFVILSAHKLVFIGDTLSRQAKAADVRSQVTHYSN<br>LLCDLLRGIVATTKAAALQYPSPSAAQDMVDRVKELGHSTQQFRRVLGQL<br>AAA |
| GST-Nck | Human (*Homo sapiens*) | MSPILGYWKIKGLVQPTRLLLEYLEEKYEEHLYERDEGDKWRNKKFELGLEF<br>PNLPYYIDGDVKLTQSMAIIRYIADKHNMLGGCPKERAEISMLEGAVLDIRY<br>GVSRIAYSKDFETLKVDFLSKLPEMLKMFEDRLCHKTYLNGDHVTHPDFML<br>YDALDVVLYMDPMCLDAFPKLVCFKKRIEAIPQIDKYLKSSKYIAWPLQGW<br>QATFGGGDHPPKSDLVPRGSENLYFQGHMAEEVVVVAKFDYVAQQEQE |

| | | |
|---|---|---|
| | | LDIKKNERLWLLDDSKSWWRVRNSMNKTGFVPSNYVERKNSARKASIVK<br>NLKDTLGIGKVKRKPSVPDSASPADDSFVDPGERLYDLNMPAYVKFNYMA<br>EREDELSLIKGTKVIVMEKSSDGWWRGSYNGQVGWFPSNYVTEEGDSPL<br>GDHVGSLSEKLAAVVNNLNTGQVLHVVQALYPFSSSNDEELNFEKGDVM<br>DVIEKPENDPEWWKARKINGMVGLVPKNYVTVMQNNPLTSGLEPSPPQ<br>SDYIRPSLTGKFAGNPWYYGKVTRHQAEMALNERGHEGDFLIRDSESSPN<br>DFSVSLKAQGKNKHFKVQLKETVYSIGQRKFSTMEELVEHYKKAPIFTSEQ<br>GEKLYLVKHLS |
| His-N-WASP | Rat (*Rattus norvegicus*) | GSEFKEKKKGKAKKKRAPPPPPPSRGGPPPPPPPPHSSGPPPP<br>PARGRGAPPPPPSRAPTAAPPPPPPSRPGVVVPPPPPNRMYPH<br>PPPALPSSAPSGPPPPPPLSMAGSTAPPPPPPPPPPPPGPPPPPG<br>LPSDGDHQVPASSGNKAALLDQIREGAQLKKVEQNSRPVSCSG<br>RDALLDQIRQGIQLKSVSDGQESTPPTPAPTSGIVGALMEVMQK<br>RSKAIHSSDEDEDDDDEEDFEDDDEWED |
| His-SUMO-FAK IDR | Mouse (*Mus musculus*) | MGSSHHHHHHSSGLVPRGSASMSDSEVNQEAKPEVKPEVKPETHINLKV<br>SDGSSEIFFKIKKTTPLRRLMEAFAKRQGKEMDSLRFLYDGIRIQADQTPED<br>LDMEDNDIIEAHREQIGGCMESRRQATVSWDSGGSDEAPPKPSRPGYPS<br>PRSSEGFYPSPQHMVQTNHYQVSGYPGSHGIPAMAGSIYQGQASLLDQT<br>ELWNHRPQEMSMWQPSVEDSAALDLRGMGQVLPPHLMEERLIRQQQE<br>MEEDQRWLEKEERFLKPDVRLSRGSIDREDGSFQGPTGNQHIYQPVGKP<br>DPAAPPKKPPRPGAPGHLSNLSSISSPADSYNEGVKLQPQEISPPPTA |
| His-SUMO-Abl IDR | Human (*Homo sapiens*) | MGSSHHHHHHHHGSGLVPRGSASMSDSEVNQEAKPEVKPEVKPETHINL<br>KVSDGSSEIFFKIKKTTPLRRLMEAFAKRQGKEMDSLRFLYDGIRIQADQTP<br>EDLDMEDNDIIEAHREQIGGRGSELPTKTRTSRRAAEHRDTTDVPEMPHS<br>KGQGESDPLDHEPAVSPLLPRKERGPPEGGLNEDERLLPKDKKTNLFSALIK<br>KKKKTAPTPPKRSSSFREMDGQPERRGAGEEEGRDISNGALAFTPLDTAD<br>PAKSPKPSNGAGVPNGALRESGGSGFRSPHLWKKSSTLTSSRLATGEEEG<br>GGSSSKRFLRSCSASCVPHGAKDTEWRSVTLPRDLQSTGRQFDSSTFGGH<br>KSEKPALPRKRAGENRSDQVTRGTVTPPPRLVKKNEEAADEVFKDIMESSP<br>GSSPPNLTPKPLRRQVTVAPASGLPHKEEAGKGSALGTPAAAEPVTPTSKA<br>GSGAPGGTSKGPAEESRVRRHKHSSESPGRDKGKLSRLKPAPPPPPAASA<br>GKAGGKPSQSPSQEAAGEAVLGAKTKATSLVDAVNSDAAKPSQPGEGLK<br>KPVLPATPKPQSAKPSGTPISPAPVPSTLPS |
| His-SUMO-mEGFP-DDX21-IDR WT | Human (*Homo sapiens*) | MGSSHHHHHHHHGSGLVPRGSASMSDSEVNQEAKPEVKPEVKPETHINL<br>KVSDGSSEIFFKIKKTTPLRRLMEAFAKRQGKEMDSLRFLYDGIRIQADQTP<br>EDLDMEDNDIIEAHREQIGGRGSMVSKGEELFTGVVPILVELDGDVNGHK<br>FSVSGEGEGDATYGKLTLKFICTTGKLPVPWPTLVTTLTYGVQCFSRYPDH<br>MKQHDFFKSAMPEGYVQERTIFFKDDGNYKTRAEVKFEGDTLVNRIELKG<br>IDFKEDGNILGHKLEYNYNSHNVYIMADKQKNGIKVNFKIRHNIEDGSVQL<br>ADHYQQNTPIGDGPVLLPDNHYLSTQSKLSKDPNEKRDHMVLLEFVTAAG<br>ITLGMDELYKGGSGGSENLYFQGGIRGPREGYGGFRGQREGSRGFRGQR<br>DGNRRFRGQREGSRGPRGQRSGGGNKSNRSQNKGQKRSFSKAFGQ |

| | | |
|---|---|---|
| His-SUMO-mEGFP-DDX21-IDR R/S | Human (*Homo sapiens*) | MGSSHHHHHHHHGSGLVPRGSASMSDSEVNQEAKPEVKPEVKPETHINLKVSDGSSEIFFKIKKTTPLRRLMEAFAKRQGKEMDSLRFLYDGIRIQADQTPEDLDMEDNDIIEAHREQIGGRGSMVSKGEELFTGVVPILVELDGDVNGHKFSVSGEGEGDATYGKLTLKFICTTGKLPVPWPTLVTTLTYGVQCFSRYPDHMKQHDFFKSAMPEGYVQERTIFFKDDGNYKTRAEVKFEGDTLVNRIELKGIDFKEDGNILGHKLEYNYNSHNVYIMADKQKNGIKVNFKIRHNIEDGSVQLADHYQQNTPIGDGPVLLPDNHYLSTQSKLSKDPNEKRDHMVLLEFVTAAGITLGMDELYKGGSGGSENLYFQGGIRGPSEGYGGFSGQSEGSSGFSGQSDGNSSFSGQSEGSSGPSGQSSGGGNKSNSSQNKGQKSSFSKAFGQ |

**Table S1. Recombinant protein sequences used in this study**

Data S1.

**Phase separation prediction scores for human kinases**. “No PS likely all predictors” tab includes kinases with a score of <0.3 for three independent predictors. “PS likely all predictors” tab includes kinases with a score of >0.7 for three independent predictors. “PS likely one predictor” tab includes kinases with a score of >0.7 for at least one predictor.

### Supplemental References


1. Schindelin, J., Arganda-Carreras, I., Frise, E., Kaynig, V., Longair, M., Pietzsch, T., Preibisch, S., Rueden, C., Saalfeld, S., Schmid, B., et al. (2012). Fiji: an open-source platform for biological-image analysis. Nat Methods *9*, 676–682. https://doi.org/10.1038/nmeth.2019.

2. Randolph, L.N., Bao, X., Zhou, C., and Lian, X. (2017). An all-in-one, Tet-On 3G inducible PiggyBac system for human pluripotent stem cells and derivatives. Sci Rep *7*, 1549. https://doi.org/10.1038/s41598-017-01684-6.

3. Niebling, S., Veith, K., Vollmer, B., Lizarrondo, J., Burastero, O., Schiller, J., Struve García, A., Lewe, P., Seuring, C., Witt, S., et al. (2022). Biophysical Screening Pipeline for Cryo-EM Grid Preparation of Membrane Proteins. Front. Mol. Biosci. *9*. https://doi.org/10.3389/fmolb.2022.882288.

4. Xie, Y., Li, H., Luo, X., Li, H., Gao, Q., Zhang, L., Teng, Y., Zhao, Q., Zuo, Z., and Ren, J. (2022). IBS 2.0: an upgraded illustrator for the visualization of biological sequences. Nucleic Acids Res *50*, W420–W426. https://doi.org/10.1093/nar/gkac373.

5. Eid, S., Turk, S., Volkamer, A., Rippmann, F., and Fulle, S. (2017). KinMap: a web-based tool for interactive navigation through human kinome data. BMC Bioinformatics *18*, 16. https://doi.org/10.1186/s12859-016-1433-7.
